# Oct4 redox sensitivity potentiates reprogramming and differentiation

**DOI:** 10.1101/2023.02.21.529404

**Authors:** Zuolian Shen, Yifan Wu, Asit Manna, Chongil Yi, Bradley R. Cairns, Kimberley J. Evason, Mahesh B. Chandrasekharan, Dean Tantin

**Affiliations:** Department of Pathology, University of Utah School of Medicine, Salt Lake City, UT 84112, USA; Huntsman Cancer Institute, University of Utah School of Medicine, Salt Lake City, UT 84112, USA; Department of Oncological Sciences, University of Utah School of Medicine, Salt Lake City, UT 84112, USA; Howard Hughes Medical Institute, University of Utah School of Medicine, Salt Lake City, UT 84112, USA; Department of Radiation Oncology, University of Utah School of Medicine, Salt Lake City, UT 84112, USA

**Keywords:** Oct4 (Pou5f1), Oct1 (Pou2f1), induced pluripotent stem cells (iPSCs), oxidative stress, ubiquitylation

## Abstract

The transcription factor Oct4/Pou5f1 is a component of the regulatory circuitry governing pluripotency and is widely used to induce pluripotency from somatic cells. Here we use domain swapping and mutagenesis to study Oct4’s reprogramming ability, identifying a redox-sensitive DNA binding domain cysteine residue (Cys48) as a key determinant of reprogramming and differentiation. Oct4 Cys48 sensitizes the protein to oxidative inhibition of DNA binding activity and promotes oxidation-mediated protein ubiquitylation. *Pou5f1^C48S^* point mutation has little effect on undifferentiated embryonic stem cells (ESCs), but upon retinoic acid (RA) treatment causes retention of Oct4 expression, deregulated gene expression and aberrant differentiation. *Pou5f1^C48S^* ESCs also form less differentiated teratomas and contribute poorly to adult somatic tissues. Finally, we describe *Pou5f1^C48S^* (*Janky*) mice, which in the homozygous condition are severely developmentally restricted after E4.5. Rare animals bypassing this restriction appear normal at birth but are sterile. Collectively, these findings uncover a novel Oct4 redox mechanism involved in both entry into and exit from pluripotency.

## Introduction

The early mammalian embryo contains undifferentiated, pluripotent cells capable of generating all embryonic tissue lineages. Cultured embryonic stem cells (ESCs) have similar capabilities (Evans and Kaufman 1981). The transcription factor Oct4 governs the establishment of undifferentiated inner cell mass and epiblast cells, as well germline cells and ESCs (Scholer et al. 1990b; Morey et al. 2015). Both ESC differentiation and blastocyst implantation are followed by Oct4 loss (Nichols et al. 1998). Together with factors such as Nanog and Sox2, Oct4 forms a network of pluripotency regulators that enforce their own and each other’s expression (Boyer et al. 2005).

Oct4 is widely used to generate induced pluripotent stem cells (iPSCs) from somatic cells (Takahashi and Yamanaka 2006). Reprogramming methods that omit Oct4 nevertheless rely on activation of the endogenous *Pou5f1* (*Oct4*) gene (Heng et al. 2010; Gao et al. 2013). Reprogramming strategies that increase efficiency (Esteban et al. 2010; Soufi et al. 2012; Vierbuchen and Wernig 2012; Costa et al. 2013; Rais et al. 2013), or improve the quality of iPSC clones (Yuan et al. 2011; Buganim et al. 2014; Chen et al. 2015; Velychko et al. 2019) reveal trade-offs: efficiently generating iPSC that are of non-optimal quality, or inefficiently generating iPSCs with improved developmental potential.

Pluripotent cells co-express Oct4 together with the widely expressed prototypic POU transcription factor Oct1/Pou2f1 (Okamoto et al. 1990; Rosner et al. 1990; Suzuki et al. 1990). The two proteins share similar DNA binding domains (DBDs) and consensus DNA binding sequences (Tantin 2013). They also associate with many of the same target genes (Ferraris et al. 2011; Perovanovic et al. 2023). Nevertheless, Oct1 and other mammalian Oct4 paralogs do not efficiently reprogram (Takahashi and Yamanaka 2006; Feng et al. 2009). In contrast, other reprogramming factors such as Sox2, Klf4 and c-Myc can be replaced with close paralogs (Nakagawa et al. 2008; Shi et al. 2008). Structure/function studies comparing Oct4 with its paralogs have identified reprogramming determinants (Esch et al. 2013; Jin et al. 2016; Jerabek et al. 2017). An example is three amino acid residues in the Oct4 DNA binding domain (DBD) but absent in Oct6/Pou3f1 needed to form Sox2 heterodimers. Introducing these residues, plus the Oct4 N- and C-termini, into Oct6 results in efficient reprogramming (Jerabek et al. 2017). Interestingly several Oct4 paralogs including Oct1 contain these residues but are still reprogramming-incompetent, indicating the presence of additional reprogramming determinants present in Oct4 (and Oct6) but absent in Oct1.

Recent studies show that omission of Oct4 from reprogramming cocktails, despite being inefficient, results in qualitatively superior iPSC clones with enhanced developmental potential (An et al. 2019; Velychko et al. 2019). The counterproductive activities of Oct4 are caused by transient promotion of gene expression programs that deviate cells from correct reprogramming trajectories (Velychko et al. 2019). A logical extension of these findings is that transient mitigation of counterproductive Oct4 activities may enable reprogramming.

Here, we identify an Oct4 DBD cysteine residue (Cys48) as a central reprogramming determinant. A serine residue is present in Oct1 at this position. In the presence of the Oct4 N-terminus, mutating Oct1’s serine to cysteine confers robust reprogramming activity. Conversely, mutating Oct4’s cysteine to serine reduces reprogramming efficiency by ∼60%. Cys48 sensitizes Oct4 DNA binding activity to oxidative stress and potentiates protein ubiquitylation. CRISPR-edited Oct4^C48S^ embryonic stem cell (ESC) lines are phenotypically normal until differentiated, following which they abnormally retain Oct4 protein expression and pluripotency gene expression signatures. Differentiating Oct4^C48S^ cells proliferate less and undergo apoptosis at higher rates compared to parental controls. Mutant ESCs also form less differentiated teratomas and contribute poorly to the development of viable adult mice. Oct4^C48S^ mutant mice (dubbed *Janky* mice, *Jky*) appear phenotypically normal and fertile in the heterozygous condition, while homozygotes are severely developmentally restricted after the blastocyst stage. Cumulatively, our findings identify an Oct4 redox sensitivity mechanism that restrains protein function, but nevertheless functions positively in reprogramming and normal differentiation.

## Results

### Oct4 vs. Oct1 reprogramming potential concentrates in a DBD cysteine

The two DNA binding subdomains of mouse Oct1 and Oct4, termed the POU-specific and POU-homeodomain (POU_S_ and POU_H_), share 71% and 59% identity, respectively (Fig. 1A). In contrast, the N- and C-termini, and the linker between the subdomains, are divergent. The human proteins are structured similarly. The high similarity allows the amino acid numbering of the DBDs to be standardized by referring to the first amino acid of POU_S_ as residue 1 (Fig. 1A). To measure reprogramming efficiency, we transduced mouse embryonic fibroblasts (MEFs) expressing GFP under the control of endogenous *Pou5f1* (Lengner et al. 2007) with lentiviruses expressing the four Yamanaka factors (Sommer et al. 2009) (Fig. 1B). To identify Oct4 determinants important for reprogramming, we replaced Oct4 within this construct with Oct1 or chimeric proteins containing the distinct DBDs and N- and C-termini (Fig. 1B). We included C-terminal FLAG-tags on all proteins to track expression. Unlike an N-terminal FLAG tag which was deleterious to Oct4 (Supplemental Fig. S1), a C-terminal FLAG tag reprograms with similar efficiency (Fig. 1C). Prior work comparing Oct4 to Oct6 identified Oct4 K40 (Fig. 1A) as important for reprogramming (Jin et al. 2016). We therefore also mutated Oct1^N40^ within POU_S_ to lysine in some constructs (N40K, Fig. 1B). Substituting Oct1 for Oct4 eliminated reprogramming potential (Fig. 1D,E), setting a baseline for domain-swaps and mutations. All of the chimeric constructs showed poor activity, with constructs containing the Oct4 N-terminus generating the most colonies (∼5% the level of WT Oct4, Fig. 1D,E). These findings are consistent with work establishing the importance of the Oct4 N-terminus (Boija et al. 2018). The raw data for Fig. 1D are shown in Supplemental Table S1. Similar data were obtained at multiple timepoints (Supplemental Fig. S2A), indicating that the results reflect qualitative rather than kinetic differences. Anti-FLAG immunoblotting showed similar expression (Fig. 1F). Viral titers from six tested vectors from this experiment were also similar (Supplemental Fig. S2B). Because a construct with both the Oct4 N- and C-termini (“4N/C”) displayed only ∼5% efficiency (Fig. 1D,E), we conclude that determinants within the Oct4 DBD confer the bulk of reprogramming potential vis-à-vis Oct1.

**Figure 1.**
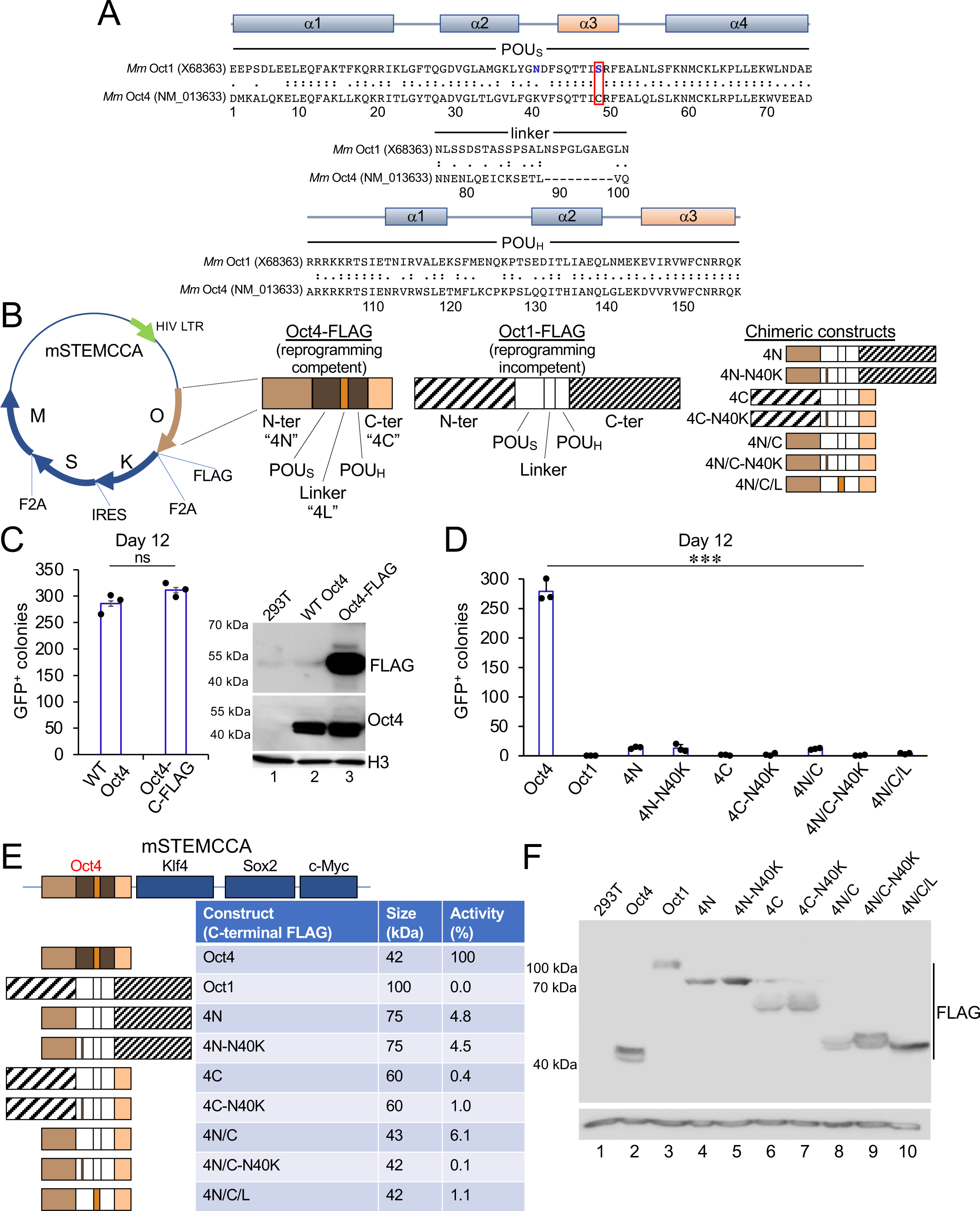
Oct4 reprogramming activity vis-à-vis Oct1 maps to the DBD. (*A*) Mouse Oct1 vs Oct4 DBD alignment. POU_S_ and POU_H_ refer to the two DNA binding sub-domains. Asparagine 40 (blue) was mutated to lysine in some constructs. The key Oct1 serine 48 residue (cysteine in Oct4) is boxed in red. (*B*) Schematic of the reprogramming vector with C-terminal Oct4 FLAG tag. HIV=human immunodeficiency virus. LTR=long terminal repeat. O=Oct4. K=Klf4. S=Sox2. M=c-Myc. The different domains of Oct4 and Oct1 are color coded. Chimeric constructs are shown at right. The construct name indicates which domain of Oct4 is included separated by slashes, e.g. 4N/C=the Oct4 N- and C-termini fused to the rest of Oct1. Point mutations are shown with a dash. (*C*) Reprogramming activity of C-terminally tagged Oct4 relative to the untagged control. Results show an average of N=3 independent experiments. Error bars show ±standard deviation. Anti-FLAG and anti-Oct4 immunoblots using tagged and untagged constructs transiently transfected into 293T cells are shown at right. (*D*) iPSC formation assay using target Oct4-GFP MEFs. Reprogramming was assessed using the number of GFP-expressing iPSC colonies. Oct4 was used as a positive control and Oct1 as a negative control. All constructs bore C-terminal FLAG-tags. N=3 experiments were performed. Error bars show ±standard deviation. (*E*) Schematic and summary of size and activity of the constructs used in D. To determine % activity, the averaged colony count using Oct4 in D was set to 100%. C-terminal FLAG-tags are not shown for simplicity. (*F*) Anti-FLAG immunoblot showing expression of the different constructs when transiently transfected into 293T cells transduced with the reprogramming vectors. Histone H3 is shown as a loading control.

In the context of the Oct4 N-terminus (“4N”), we mutagenized DBD amino acid residues that differ between Oct1 and Oct4, focusing on post-translationally modifiable residues: Lys, Ser, Thr and Cys. This analysis identified 14 different amino acids, not including Lys40 (Fig. 1A). Oct6 is capable of reprogramming when modified to enable Sox2 dimerization (Kim et al. 2020). Filtering these residues using conservation between Oct4 and Oct6 pinpointed six possible residues. Replacing one of these, Oct1 Ser48, with the Oct4 cysteine residue found at this position increased activity >7-fold over the “4N” construct containing the Oct4 N-terminus alone (Fig. 2A,B, “1-4All”). The raw data for Fig. 2A are shown in Supplemental Table S2. iPSC colonies formed under these conditions showed a pluripotent morphology and expressed GFP from the endogenous *Pou5f1* locus at levels comparable to Oct4 (Supplemental Fig. S3). The remaining activity was concentrated in the linker domain, as a construct containing the Oct4 N-terminus, Cys48 and the Oct4 linker (4N/L-S48C) had 100% activity (Fig. 2A,B). The Oct4 linker has been shown to regulate reprogramming via selective interactions with different cofactors (Esch et al. 2013; Han et al. 2022). The differences were maintained over time, with the 1-4All construct consistently generating ∼60% activity over multiple days, indicating that the differences were not kinetic in nature (Supplemental Fig. S4A). Similar results were obtained using normal primary MEFs and assessing alkaline phosphatase-positive rather than GFP-positive iPSCs (Supplemental Fig. S4B-C).

**Figure 2.**
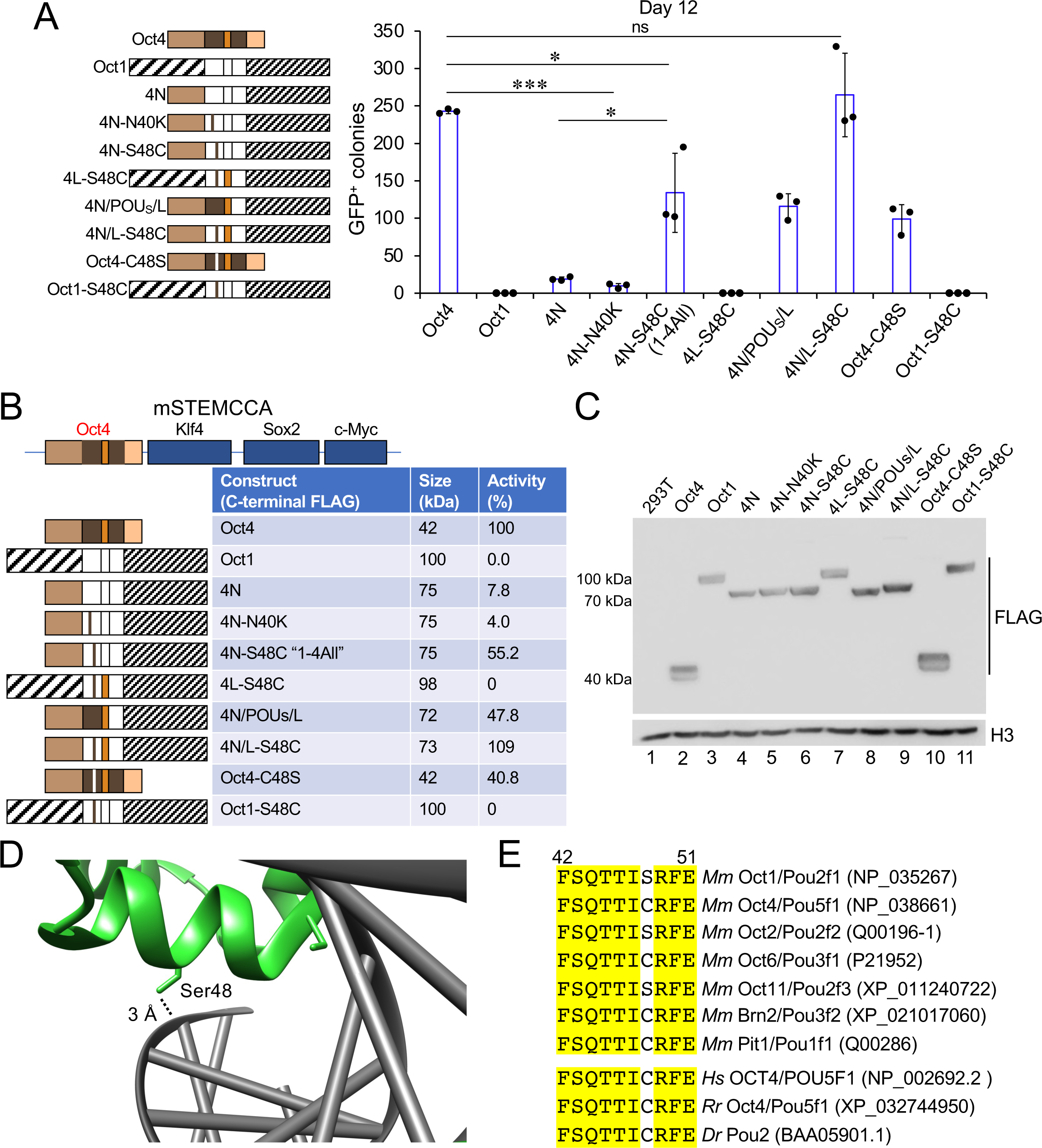
Activity of Oct4 cysteine 48 in reprogramming. (*A*) iPSC formation assay using Oct1-Oct4 lentiviral constructs that manipulate cysteine 4. Oct4 was used as a positive control and Oct1 as a negative control. All constructs bore C-terminal FLAG-tags. N=3 experiments. Error bars show ±standard deviation. (*B*) Schematic and summary of size and activity of the constructs used in A. The activity of Oct4 in A was set to 100%. (*C*) Anti-FLAG immunoblot showing expression of the different constructs when transiently transfected into 293T cells transduced with the reprogramming vectors. Histone H3 is shown as a loading control. (*D*) Detail from an Oct1:octamer DNA co-crystal structure (PDB ID: 1OCT)(Klemm et al. 1994) showing Ser48 making hydrogen bond contacts with the DNA backbone. (*E*) Paralog comparison of mouse POU protein amino acid residues corresponding to α-helix 3 of the POU_S_ sub-domain.

An Oct4^C48S^ construct reduced activity by 60% (Fig. 2A,B), indicating that over half of Oct4’s differential reprogramming activity vis-à-vis Oct1 operates through this residue. The reciprocal Oct1^S48C^ mutation did not reprogram (Fig. 2A,B, Oct1-S48C), indicating that Cys48 is not sufficient, and highlighting its combined importance with the Oct4 N-terminus for reprogramming. The proteins were equivalently expressed (Fig. 2C). These results indicate that in the context of the Oct4 N-terminus, Cys48 confers robust reprogramming activity. In an Oct1 DBD:DNA structure (Klemm et al. 1994), Ser48 hydrogen bonds with the DNA backbone (Fig. 2D). In an Oct4:Sox2:nucleosome structure (Michael et al. 2020), the Cys48 sulfhydryl also allows for hydrogen bonding. Predictably, Cys48 mutation to glycine greatly diminishes Oct4 DNA binding in vitro (Marsboom et al. 2016). Oct6 also contains a cysteine at this position, as does human Oct4 (Fig. 2E). The *Danio rerio* Oct1/4 paralog Pou2 also contains a cysteine at this position (Fig.2E), suggesting that cysteine may be the ancestral residue. These structural studies and our findings indicate that Ser48 and Cys48 have minimal differences in DNA binding, but that Cys48 nevertheless plays key roles in reprogramming.

### Oct4 Cys48 mediates degradation in response to oxidative stress

Oct4 is hypersensitive to oxidative stress (Lickteig et al. 1996; Guo et al. 2004; Marsboom et al. 2016) in a manner reversed by thioredoxin (Guo et al. 2004), implicating cysteine thiols in redox sensitivity. We reproduced this result using electrophoretic mobility shift assays (EMSA) and DNA containing a canonical binding element (Supplemental Fig. S5), suggesting that Cys48 may potentiate reprogramming through a redox mechanism. We used lysates from 293T cells transfected with WT or C48S Oct4 to measure DNA binding. To measure redox sensitivity, we pre-treated Oct4 with the oxidizing agent diamide prior to addition of DNA. 293T cells express endogenous Oct1 as an internal control (Fig. 3A, lanes 2-3). Oct4 bound to octamer DNA specifically (Fig. 3A, lanes 7-8) and was hypersensitive to diamide treatment (lanes 9-11). Oct4^C48S^ bound DNA equivalently (lane 13) but was comparatively resistant to oxidation (lanes 14-16). In contrast, little sensitivity was observed with Oct1. Quantification from three replicates is shown in Fig. 3B. The recombinant Oct4 contained a C-terminal FLAG tag, allowing for purification (Supplemental Fig. S6). Similar relative resistance to oxidative inhibition of DNA binding was observed using purified recombinant C48S Oct4 (Fig. 3C). Quantification from three replicates is shown in Fig. 3D. These results indicate that Oct4 Cys48 mediates oxidative inhibition of DNA binding.

**Figure 3.**
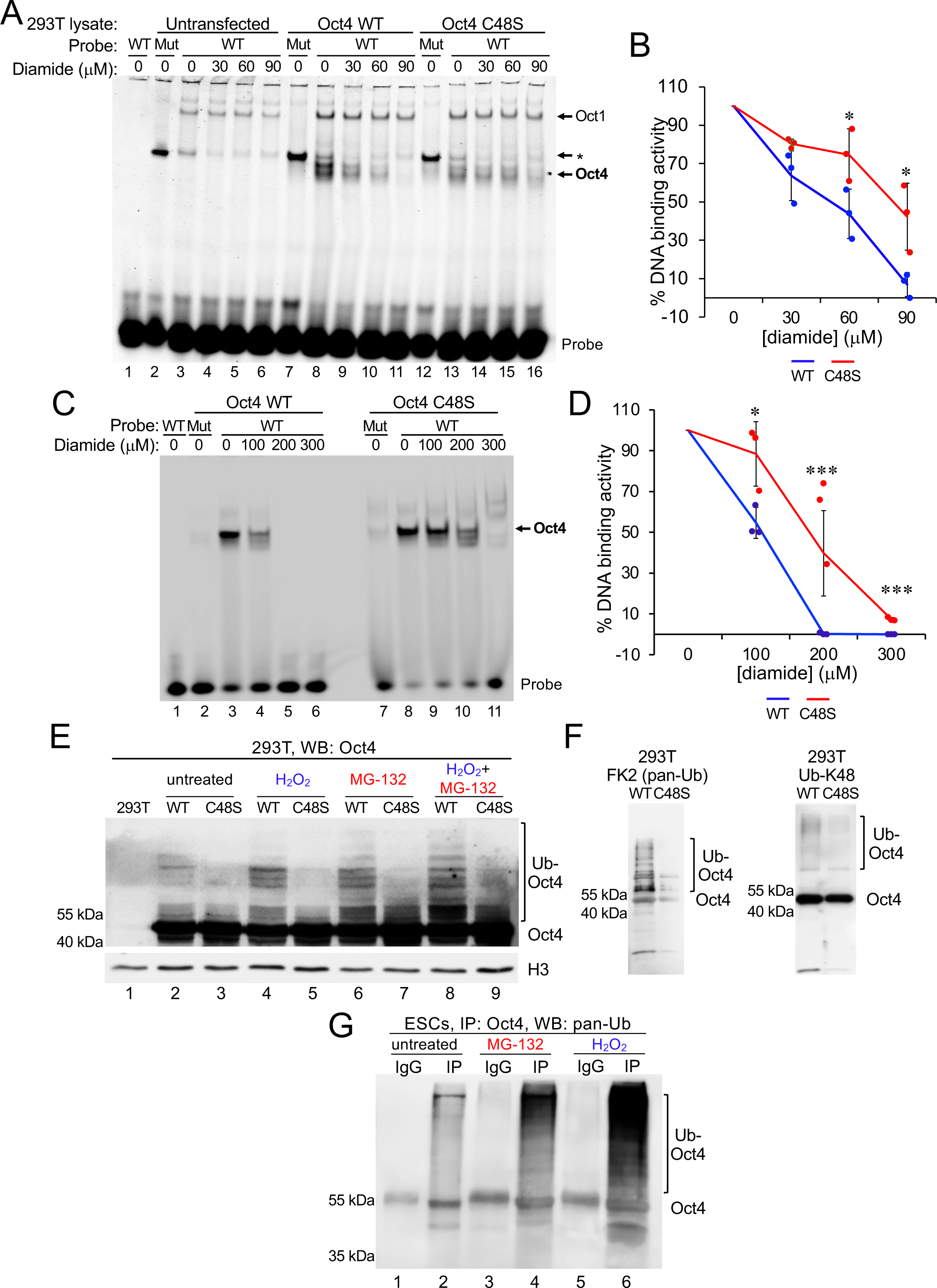
Oct4 cysteine 48 confers DNA sensitivity to oxidative stress and oxidative stress-induced ubiquitylation. (*A*) EMSA using canonical octamer DNA and wild type of C48S full-length Oct4. 293T cells were transiently transfected with plasmids encoding C-terminal FLAG-tagged Oct4 constructs. Protein lysates were made and immediately used for EMSA with increasing amounts of diamide. The larger redox-resistant Oct1 protein that binds wild type (WT) but not mutant (Mut) probe is highlighted with an arrow. Asterisk shows a redox sensitive band with superior binding to mutant rather than octamer DNA. (*B*) Quantification of Oct4 band intensity relative to no diamide, which was set to 100%. N=3 independent experiments. Error bars show ±standard deviation. (*C*) Oct4 containing a C-terminal streptavidin tag was purified from 293T cell lysates using anti-FLAG beads. Isolated protein was used in assays similar to A. (*D*) Quantification of C and two similar experiments. Error bars show ±standard deviation. (*E*) Highly exposed Oct4 immunoblot using lysates prepared from 293T cells transfected with constructs expressing C-terminally FLAG/Twin-Strep-tagged Oct4. Subsets of cells were pre-treated with H_2_O_2_ (1 mM for 2 hr), MG-132 (10 μM for 2 hr) or both. All samples contained MG-132 in the lysis buffer to prevent degradation. Histone H3 is shown as a loading control. (*F*) Purified wild type and C48S Oct4 was immunoblotted using antibodies against specific ubiquitin linkages. (*G*) Oct4 and control immunoprecipitates from wild type ESCs were immunoblotted using FK2 antibodies. Cells were treated with 10 μM MG-132 for 2 hr or 2.5 mM H_2_O_2_ for 2 hr.

Oct4 oxidation has been linked to its degradation (Marsboom et al. 2016). We noted in over-exposed immunoblots the presence of multiple slow-migrating high molecular weight forms of wild type Oct4 which were attenuated with C48S mutation (Fig. 3E, lanes 2-3). Hydrogen peroxide treatment augmented the banding pattern using wild type but not mutant protein (lanes 4-5). The bands were also increased using MG-132, consistent with ubiquitylated species (lanes 6-7). H_2_O_2_ and MG-132 co-treatment further increased the pattern (lanes 8-9). None of these treatments affected short-term cell viability (Supplemental Fig. S7).

We immunoblotted purified Oct4 with antibodies specific to different ubiquitin species. Antibodies recognizing both mono- and poly-ubiquitylated proteins revealed strong diminution with Oct4^C48S^ as expected (Fig. 3F, FK2). Diminution was observed with K48 linkage-specific polyubiquitin antibodies (Fig. 3F), while K63 linkage-specific antibodies showed modest differences (Supplemental Fig. S8). Similar ubiquitylated species were observed using endogenous Oct4 in wild type ESCs (Fig. 3G, lane 2) that were augmented by H_2_O_2_ and MG-132 (lanes 4 and 6). We conclude that Cys48 promotes Oct4 ubiquitylation and degradation in response to oxidative stress, at least in part via catalysis of K48- linked polyubiquitin chains.

### Oct4^C48S^ ESCs differentiate abnormally

We generated ESCs with a single-base missense mutation in the endogenous *Oct4* (*Pou5f1*) locus that converts Cys48 to Ser. Two silent point mutations were also engineered to create an *Xba*I site to monitor targeting and facilitate genotyping. Three low-passage parent ESC lines of different strain backgrounds were chosen for targeting. Multiple independent clones were generated for each line. Homozygous mutation was confirmed by high-resolution melting analysis and resequencing (Supplemental Fig. S9A and Fig. 4A). Tested clones were karyotypically normal and similar to parent ESCs, with the proportion of diploid cells ranging between 71 and 93%. Undifferentiated mutant ESC clones expressed Oct4, Sox2 and Nanog at similar levels and were morphologically indistinguishable from their parent clones (Fig.4B-D, Supplemental Fig. S9B-D).

**Figure 4.**
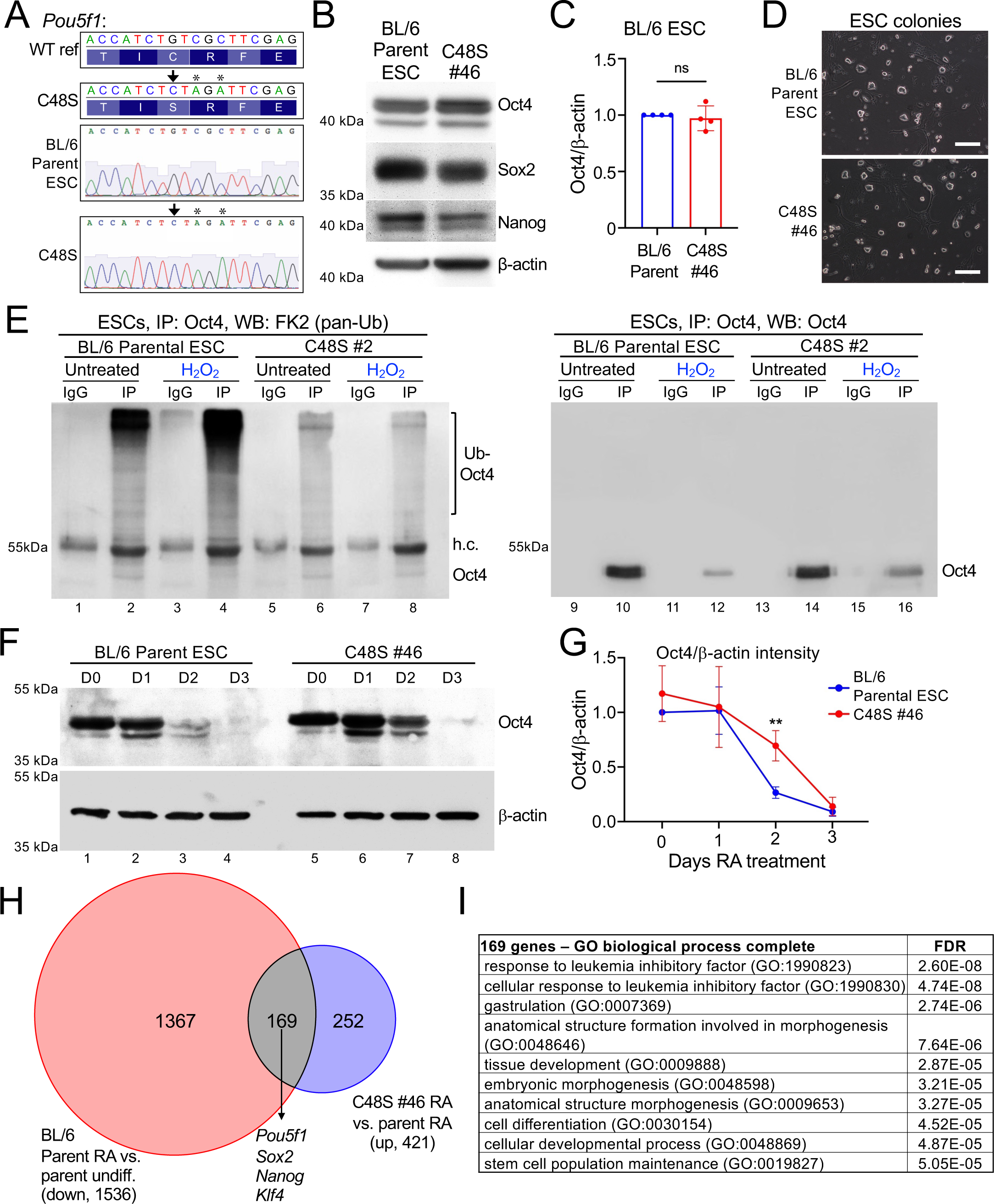
Oct4^C48S^ ESCs are phenotypically normal but retain Oct4 expression when differentiated. (*A*) *Top:* Schematic for CRISPR-mediated *Pou5f1* (*Oct4*) mutation. A single nucleotide change (arrow) converts the wild type cysteine to the serine present in Oct1. Asterisks depict two silent point mutations present in the guide RNA to monitor mutagenesis. *Bottom:* DNA sequencing trace of homozygous mutant clone. (*B*) An example C57BL/6 ESC clone (#46) was immunoblotted for Oct4, Sox2, Nanog and β-actin as a loading control. (*C*) Quantification of Oct4 protein levels from N=4 immunoblots. Levels were normalized to β-actin. Error bars depict ±standard deviation. (*D*) Example images of pluripotent colonies from a parental ESC line and derived Oct4^C48S^ ESC line (#46). Images were collected at 10× magnification. Scale bar = 0.2 mm. (*E*) Pan-Ub and control Oct4 immunoblots of Oct4 immunoprecipitates from parent and Oct4^C48S^ (clone #2) ESC lines. Cells were grown on gelatin without feeders. Normal culture medium was replaced with medium lacking supplemental glutamine and β- mercaptoethanol 2 hr prior to the experiment. Where indicated, cells were treated with 2.5 mM H_2_O_2_ for a further 2 hr. Oct4 was immunoprecipitated using a rabbit polyclonal antibody (Abcam) and immunoblotted using monoclonal mouse antibodies against pan-Ub and Oct4. (*F*) Parent or derived Oct4^C48S^ ESCs were differentiated using RA. Lysates were collected at the indicated times and probed for Oct4 by immunoblotting. β-actin was used as a loading control. (*G*) Average quantified Oct4/β-actin levels from N=3 immunoblots conducted with mutant ESC clone #46. Error bars depict ±standard deviation. (*H*) mRNA from parental and #46 mutant cells differentiated for 2 days with RA, and undifferentiated controls, were transcriptionally profiled using RNA-seq. Significantly increased genes in differentiating C48S relative to parent cells (421 genes) were intersected with the set of genes down regulated in parental cells upon differentiation (1536 genes), to yield 169 genes. (*I*) PANTHER GO terms associated with 169 genes from G identified using the “biological process” ontology.

To test if C48S mutation protects endogenous Oct4 against oxidative stress, we studied Oct4 ubiquitination in parent and mutant ESCs. Oct4 was immunoprecipitated from ESC lysates, and the resulting material was immunoblotted using pan-Ub (FK2) and Oct4 antibodies. Ubiquitylation of endogenous Oct4 in ESCs was clearly visualized and increased with H_2_O_2_ exposure (Fig. 4E, lanes 1-4), while Oct4^C48S^ ubiquitylation was attenuated (lanes 5-6) and unaffected by oxidative stress (lanes 7-8). Oct4 immunoblotting confirmed that the protein was immunoprecipitated, with decreased Oct4 recovered with oxidative stress in parental cells and more Oct4 recovered in mutant cells (Fig. 4E, right side).

Cells were then differentiated in vitro using retinoic acid (RA) to assess Oct4 expression, morphology and cell counts. Oct4 protein expression was abnormally retained early during differentiation (Fig. 4F, lanes 3 and 7). Quantification from multiple experiments revealed that the effect was transient, with a >2.8-fold increase of Oct4 at day 2 that normalized by day 3 (Fig. 4G). Similar results were obtained using the same clone compared to parent cells electroporated without CRISPR RNP (Supplemental Fig. S9E). These results were recapitulated using a second clone from the same parental ESC line (Supplemental Fig. S9F), as well as a clone from a different 129×C57BL/6 ESC line, which differentiated more slowly and showed greater differences (Supplemental Fig. S9G).

We performed transcriptional profiling of parent and mutant cells at RA differentiation day 2, as well as undifferentiated controls. Three replicates per condition were used, which showed tight concordance (Supplemental Fig. S10A). 20-31 million sequence reads were generated for each sample, ∼80% of which aligned to the *mm39* reference genome. We identified 421 genes with significantly elevated gene expression in differentiating mutant relative to parent cells (*p*<0.001, >2.5-fold, Fig. 4H, Supplemental Table S3). Intersecting these genes with the set of genes normally decreased upon differentiation of parent cells at this timepoint (1536 genes) showed that 40% of the 421 genes with retained expression in mutant cells (169 genes) are normally down-regulated with differentiation. These included *Oct4* itself (*Pou5f1*), as well as *Sox2*, *Nanog* and *Klf4* (Fig. 4H). Comparing these genes with established Oct4 ChIP-seq (King and Klose 2017) identified 114 of 169 genes (67%) as direct Oct4 targets (Supplemental Table S3). The complete set of 169 genes were enriched for gene ontology (GO) terms associated with pluripotency such as “response to leukemia inhibitory factor” (including *Pou5f1*, *Klf4*, and *Fgf4*) and “stem cell population maintenance” (including *Nanog*, *Pou5f1*, *Sox2* and *Fgf4*, Fig. 4I). Similar results were obtained using a hierarchical clustering approach. We utilized the 100 most differentially expressed genes upon differentiation for both genotypes (150 total genes). Although the differentiating parental and mutant replicates clustered together, there was a cohort of pluripotency associated genes (including *Pou5f1*) whose down-regulation was blunted in mutant cells (Supplemental Fig. S10B). A second cohort of differentiation-specific genes showed poorer expression in mutant cells upon differentiation. These included *Krt18*, *Mest*/*Peg1* and *Dag1* (Supplemental Fig. S10B). Comparing the complete set of 150 genes to Oct4 ChIP-seq (King and Klose 2017) identified 117 (78%) as direct Oct4 targets (Supplemental Table S3). These results indicate that Oct4^C48S^ ESCs inappropriately delay loss of pluripotency gene expression signatures during differentiation.

The level and duration of retained Oct4 expression during differentiation can affect lineage specification (Wang et al. 2012; Aksoy et al. 2013; DeVeale et al. 2013; Radzisheuskaya et al. 2013). To determine the effects of Oct4^C48S^ mutation on differentiation, we visualized cells treated with RA microscopically. These experiments revealed a significant depletion of cells beginning at RA differentiation day 3 (Fig. 5A,B, D6-D12). The decreased cell numbers could be due to decreased proliferation and/or increased cell death. We studied proliferation using cell trace violet (CTV) and apoptosis using TUNEL staining. Adherent, viable differentiating C48S cells showed decreased proliferation at days 3 and 4 (Fig. 5C) and increased apoptosis at day 2 (Fig. 5D). ESCs differentiating with RA also experience an increase in global reactive oxygen species levels at day 3, though specific oxidizing molecules at specific subcellular locations may be higher (Fig. 5E). Finally, we used mutant ESCs to form embryoid bodies (EBs) and subsequently generate cardiomyocytes. Differentiated cardiomyocytes form gap junctions, with functionality scorable by spontaneous and synchronous beating of groups of cells (Lynch et al. 2018). We found that unlike parent cells which formed beating cardiomyocytes robustly, two different mutant ESC lines completely failed to form beating cells when differentiated (Fig. 5F and Supplemental Movies S1-S6). Beating was not rescued by extended culture, indicating that the phenotype was not kinetic in nature (not shown). These results define a functional role for Oct4 Cys48 in ESC differentiation.

**Figure 5.**
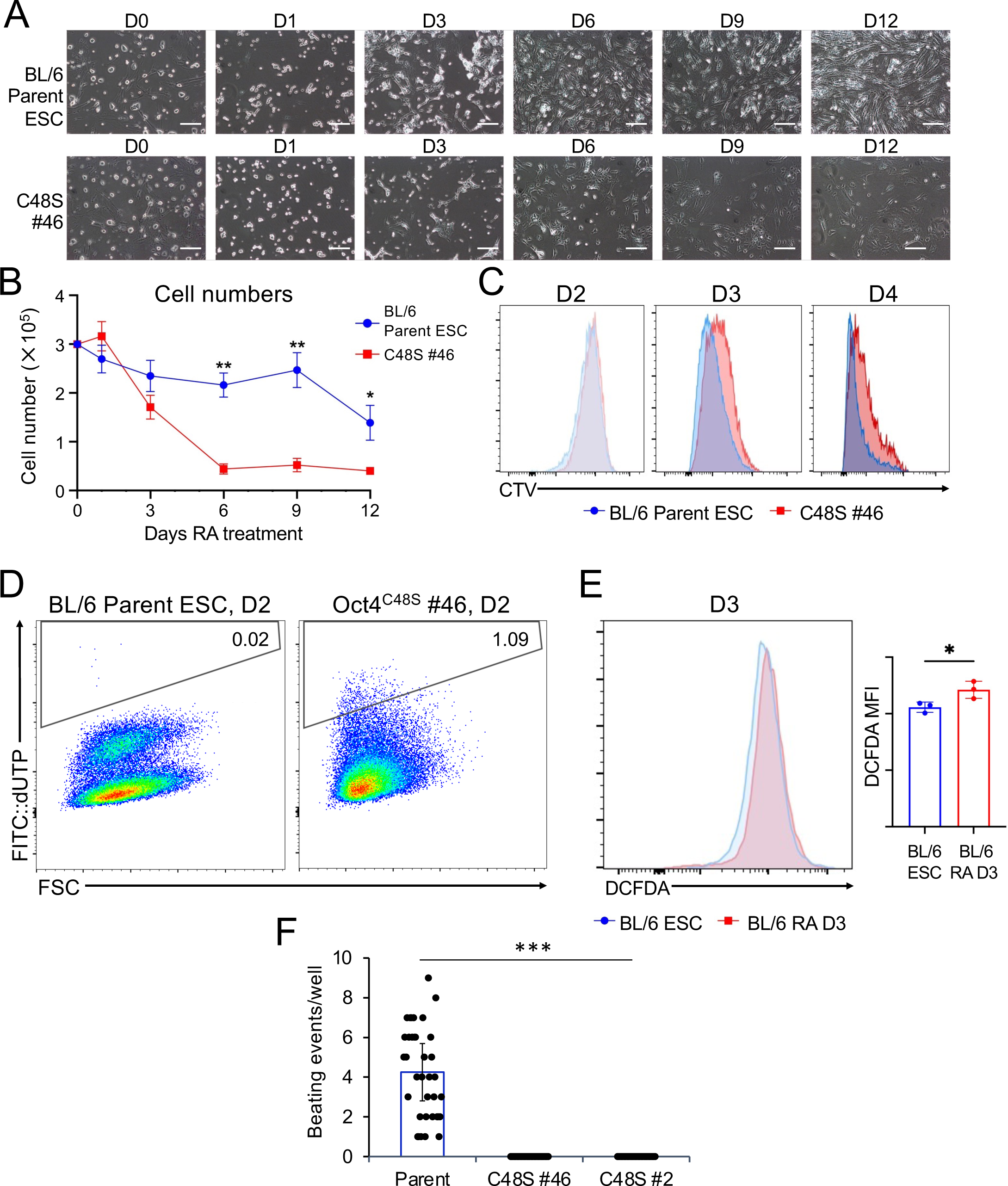
Oct4^C48S^ ESCs differentiate abnormally. (*A*) Phase-contrast images of parent and derived Oct4^C48S^ ESCs (clone #46) differentiated using RA for 12 days. Images were collected at the indicated times at 10× magnification. Scale bar = 0.2 mm. (*B*) Differentiating cells were trypsinized and counted using trypan blue exclusion using a hemacytometer. Cell counts (N=3) were averaged and plotted. Error bars depict ±standard deviation. (*C*) Proliferation was measured in parent and C48S ESCs differentiating with RA at days 2, 3 and 4. Cells were loaded with cell trace violet (CTV) at day 2. (*D*) TUNEL assay showing Oct4^C48S^ ESCs differentiated using RA. Cells were differentiated for 2 days and incubated with FITC-labeled dUTP for 1 hour at 37℃ prior to flow cytometry. (*E*) Undifferentiated and 3-day differentiating parent ESCs were incubated with DCFDA to assess ROS levels. (*F*) Parent and derived Oct4^C48S^ ESCs (clones #2 and 46) were differentiated into EBs, mesoderm lineage cells and cardiomyocytes to assess beating. Average number of beating events per well is shown for N=36 wells, with 10 EBs plated per well. Error bars depict ±standard deviation.

### Oct4^C48S^ ESCs form less differentiated teratomas and contribute poorly to embryonic development

We used mutant and control ESC lines to form teratomas in mice, which were monitored for two weeks before tumor excision. Tumors were fixed, H&E stained, and pathologically scored in a blinded fashion. Parent tumors contained areas of maturing structures such as squamous tissue, respiratory epithelium (cilia and/or goblet cells), maturing neural components, gastric foveolar epithelium and fat. In contrast, tumors formed with Oct4^C48S^ ESCs injected into the contralateral flanks of the same mice were more homogeneous with few maturing components, instead harboring immature neuroepithelial elements, immature cartilage, embryonal carcinoma and yolk sac (Fig. 6A,B). Additional images are shown in Supplemental Fig. S11. Similar results were obtained using a wild type and mutant teratomas derived from a different ESC line (Fig. 6C). At later time points, larger mutant teratomas began to accumulate mature structures and the differences largely normalized (Supplemental Fig. S12).

**Figure 6.**
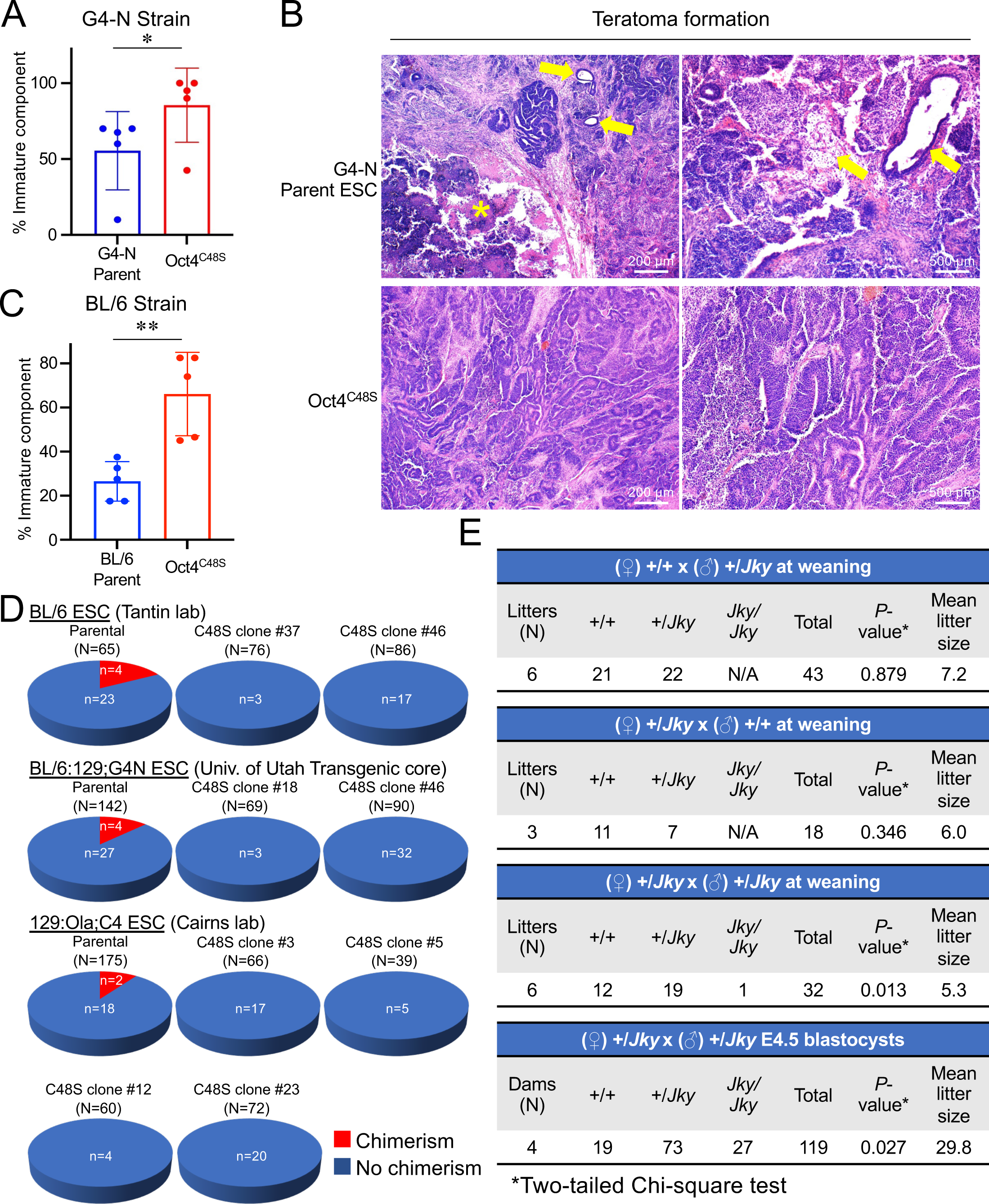
Oct4^C48S^ ESCs form teratomas lacking maturing structures and contribute poorly to adult chimeras. (*A*) 1×10^6^ parental or Oct4^C48S^ C57BL/6:129 ESCs (clone G4-N) were injected into contralateral flanks or NRG mice. After 15 days, tumors were harvested, fixed, sectioned, stained with H&E, deidentified and scored pathologically. The percent of cells within the tumor containing undifferentiated components was averaged and plotted. N=5 independent tumors for each condition combined from two separate experiments. For each tumor, values represent the average from a pair of slides. Two nonadjacent sections from each tumor were used. Error bars depict ±standard deviation. *P*-value was generated using an unpaired one-tailed T-test. (*B*) Representative images are shown. Mature (differentiated) components, e.g. airway epithelium in teratomas derived from parental ESCs are highlighted by yellow arrows. Asterisk indicates area of necrosis. (*C*) Similar data as in (A), except generated using an independently derived mutant from a different ESC line. N=5 independent tumors. (*D*) Three different parent ESC lines and 2-4 derived lines of CRISPR-mutated Oct4^C48S^ ESCs were injected into recipient blastocysts (N=number injected) and implanted into pseudo-pregnant animals. For each line, the proportion of chimeric offspring at weaning is shown as a pie chart. All karyotypes were confirmed to be similar to parent lines (between 72% and 92% normal).

One hallmark of pluripotency is the ability to contribute to cells and tissues of the embryo (De Los Angeles et al. 2015). We conducted chimerism assays to assess the ability of Oct4^C48S^ ESCs to contribute to development. We used eight lines of mutant ESCs, derived from three parent lines of different strain backgrounds to minimize background and clonal effects. Compared to the parent cells, which contributed to offspring at a rate of 10-15% (Fig. 6C), none of the derived Oct4^C48S^ ESCs were able to contribute to adult mice (Fig. 6D). The results indicates that Oct4^C48S^ ESCs contribute poorly to embryonic development. These results identify a role for Oct4 redox sensitivity in differentiation and development.

### Oct4^C48S^ homozygous mutant mice show developmental lethality after the blastocyst stage

We generated germline mutant mouse lines with the Oct4^C48S^ mutation (dubbed *Janky* mice) using CRISPR modification of the endogenous *Pou5f1* gene. A heterozygous male founder was generated, which transmitted the *Jky* allele normally, as did its male and female offspring (Fig. 6E). In contrast, heterozygote intercrosses were associated with decreased litter sizes and a significant deviation from 1:2:1 Mendelian ratios, with homozygous *Jky* offspring severely attenuated (Fig. 6E). One *Jky* homozygous male was born to these crosses, which was phenotypically indistinguishable from its littermates at weaning but was sterile with underdeveloped testes (Supplemental Fig. S13A). Microscopic examination revealed significant heterogeneity in the size and diameter of the seminiferous tubules (Supplemental Fig. S13B). Immunofluorescence using spermatogonia and spermatocyte markers showed that a minority of seminiferous tubules had normal physiology (Supplemental Fig. S13C, Box 1), but many displayed abnormal localization of spermatocytes and germ cell loss. Interestingly, the DNA damage marker ψ-H2AX, which is normally associated with meiotic sex chromosome inactivation in pachytene spermatocytes (Fernandez-Capetillo et al. 2003), was not detected in *Jky*/*Jky* LIN28A^+^ spermatogonia, suggesting that loss of Oct4 redox sensing may regulate germ cell development but not spermatogonia maintenance. The developmental lethality associated with most embryos occurred after embryonic day 4.5, as blastocysts collected at this time genotyped as *Jky*/*Jky* at normal ratios (Fig. 6E). Collectively, these results identify Oct4 Cys48 as redox-sensitive and uncover dual roles for this residue in reprogramming and development.

## Discussion

Oct4 proteins levels must be precisely controlled for efficient iPSC reprogramming (Ilia et al. 2023) and exit from pluripotency (Wang et al. 2012; Aksoy et al. 2013; DeVeale et al. 2013). Nevertheless, the mechanisms by which Oct4 is post-transcriptionally regulated are poorly defined. Here, we pinpoint Oct4 redox sensitivity as an important determinant of both reprogramming and differentiation. Pluripotent cells co-express Oct4 and Oct1 (Shen et al. 2017). Despite having similar DBDs and recognizing the same 8-bp consensus octamer sequence (Tantin 2013), Oct1 and other Oct4 paralogs do not function in reprogramming (Takahashi and Yamanaka 2006). Structure/function studies comparing Oct4 with these paralogs have identified different Oct4 reprogramming determinants (Esch et al. 2013; Jin et al. 2016; Jerabek et al. 2017; Boija et al. 2018). We identify Cys48 of the Oct4 DBD as a critical reprogramming determinant that sensitizes the protein to oxidative inhibition and degradation. In Oct1, this residue is a serine. Both residues hydrogen bond with the DNA phosphate backbone.

Transcription factors with both positive and negative roles in reprogramming include Oct4 and Smad3 (Ruetz et al. 2017; An et al. 2019; Velychko et al. 2019). Recent studies show that removal of Oct4 from reprogramming cocktails is inefficient but results in superior iPSC clones with enhanced developmental potential (An et al. 2019; Velychko et al. 2019). The counter-productive activity of Oct4 is due to promotion of off-target, ectoderm-specific gene expression (Velychko et al. 2019). These findings suggest that transient down-regulation of counterproductive Oct4 activities may be an important step for efficient reprogramming. Our work suggests that Oct4 redox sensitivity fulfills such a role. The findings may provide a route to generating superior iPSC reprogramming outcomes with higher efficiency. Oct4 is hypersensitive to oxidative stress (Lickteig et al. 1996; Guo et al. 2004; Marsboom et al. 2016) in a manner reversed by thioredoxin (Guo et al. 2004). Relative to other Oct proteins, Oct4 has unusually high cysteine content in the DBD, with two cysteines in each subdomain. Cys142 (Cys50 of the POU-homeodomain) lies in the recognition helix of POU_H_ and is the distinguishing feature of POU-homeodomains (Tantin 2013). Cys48 is located within the DNA recognition helix of the other DNA binding subdomain, POU_S_. The hydrogen bond contact between Cys48 and the DNA predicts that cysteine oxidation will block DNA binding. Consistently, Oct4^C48S^ mutation maintains DNA binding and confers resistance to oxidation-induced DNA binding inhibition in vitro. The same mutation reduces reprogramming efficiency by more than half. Cysteine redox sensitivity is known to be enhanced by adjacent basic amino acid residues (Britto et al. 2002), and Cys48 is followed by a conserved Arg. Oct1 S48C mutation is sufficient to confer reprogramming activity, but only in the context of the Oct4 N-terminus. The Oct4 N-terminus is known to mediate liquid-liquid phase transitions to control target gene transcription (Boija et al. 2018).

Prior studies have identified a link between Oct4 oxidation and degradation (Marsboom et al. 2016). Cys48 strongly promotes protein ubiquitylation, in particular following exposure to oxidative stress or MG-132. Oct4 is known to be polyubiquitylated by K63-linked chains via the action of the Stub1 E3 Ub ligase (Mamun et al. 2022), however we found that K48-specific linkages were more strongly affected with Oct4^C48S^ mutation, suggesting a distinct pathway.

Endogenous Oct4^C48S^ mutation only minimally affects undifferentiated ESCs, but results in abnormalities upon RA-mediated differentiation, including retention of Oct4 expression, deregulated gene expression, decreased proliferation and increased apoptosis. Prominent roles for Oct4 in cell survival and proliferation during post-implantation embryonic development have been documented (DeVeale et al. 2013). When injected into adult mice, ESCs form teratomas containing differentiating structures representing all three germ layers. When injected into blastocysts, they contribute to adult mouse tissues (De Los Angeles et al. 2015). We found that both features become defective with Cys48 mutation. Despite being gain-of-function, mice with this germline mutation (dubbed *Janky* mice) have no observed defects in the heterozygous condition. This may be due to a necessity for increased Oct4 activity above a certain threshold to trigger a phenotype. Homozygous *Jky* mutant mice undergo developmental lethality between E4.5 and birth. Given the timing of Oct4 expression during development, we predict that defects arise in these embryos at the blastocyst and/or peri-implantation stages, though this still awaits testing. Rare *Jky* mice obtained from these crosses appear normal at weaning, suggesting that mice circumventing this developmental restriction subsequently develop normally. However, the single male mouse we obtained from these crosses was sterile. These findings are compatible with known roles for Oct4 in the germline (Scholer et al. 1990a; Kehler et al. 2004). The abnormal seminiferous tubule architecture of this mouse resembles the Pou5f1 TNAP-Cre conditional knockout (Kehler et al. 2004), suggesting a potentially central role for Oct4 redox regulation in the survival and migration of primordial germ cells. These results indicate that Cys48 is as important for differentiation and exit of pluripotency as it is for reprogramming and indicate a physiological role for Oct4 redox regulation through Cys48 during development.

## Materials and methods

### Cell culture

ESCs/iPSCs were cultured in Dulbecco’s modified Eagle medium (Sigma) with “2iV” supplementation as described previously (Shakya et al. 2015; Shen et al. 2018). Media contained 15% fetal calf serum (FCS, Avantor), 2 mM L-glutamine (ThermoFisher), 50 μM β-mercaptoethanol (2ME, Sigma), 1,000 U/mL leukemia inhibitor factor (LIF, Chemicon), 0.1 mM minimal essential medium nonessential amino acids (NEAA, ThermoFisher), 1 mM sodium pyruvate (ThermoFisher), the MEK inhibitor PD03259010 (1 μM, LC laboratories) and the glycogen synthase kinase-3β (GSK3β) inhibitor CHIR99021 (3 μM, LC Laboratories). Culture medium was also supplemented with 50 μg/mL ascorbic acid (Sigma). H_2_O_2_ (Sigma) was supplied at 1 mM for 2 hr and MG-132 (Sigma) at 10 μM for 2 hr. RA differentiation media omitted LIF, MEK inhibitor and GSK3β inhibitor, and contained retinoic acid (Sigma) at 1.5 μg/mL. Cardiomyocytes were generated using established protocols in which EBs are first generated via hanging drops and transferred to 12-well plates with differentiation medium (DMEM, 15% FCS, 2 mM L-glutamine, 0.1 mM NEAA, 1 mM sodium pyruvate, 0.1 mM 2ME, 50 μg/mL ascorbic acid) (Lynch et al. 2018). After 8 days in differentiation culture, cells were observed using an Olympus IX-51 inverted phase-contrast microscope. CTV proliferation assays were conducted using a kit (CTV Proliferation Kit for Flow Cytometry, ThermoFisher). TUNEL assays were conducted using a kit (Apo-Direct, Becton-Dickinson). Alkaline phosphatase staining was performed using a kit (System Biosciences) according to the manufacturer instructions. DCFDA staining was performed using a kit (AbCam), following the manufacturer protocol.

### FLAG-tagged lentiviral constructs for iPSC generation

The mSTEMCCA lentiviral vector expressing Oct4, Klf4, Sox2 and c-Myc was used for these studies (Somers et al. 2010). The Oct4 cDNA cassette was replaced with c-terminally FLAG-tagged full-length mouse Oct4, mouse Oct1 or chimeric constructs. FLAG-tagged Oct4 was generated by overlap PCR using primers containing the DYKDDDDK FLAG peptide sequence: (Fwd: 5’-GATTACAAGGATGACGACGATAAGGGAAGTGGCGTGAAACAGA; Rev: 5’-CTTATCGTCGTCATCCTTGTAATCGTTTGAATGCATGGGAGAG). The first fragment was amplified using a Fwd primer containing a *Not*I restriction enzyme site (5’-AATGAAAAAAGCGGCCGCCATGGCTGGACACCT) and overlapping Rev primer containing the FLAG sequence. A second fragment was amplified using the Rev primer containing an *Nde*I restriction site (5’-GGGAATTCCATATGTGTGGCCATATTATCATCGTGT) and overlapping Fwd primer. The first and second fragments were mixed as template and amplified Fed and Rev primers containing *NotI* and *Nde*I. After purification and digestion with *Not*I and *Nde*I, the fragment was ligated with the mSTEMCCA vector digested by *Not*I and *Nde*I. Similar strategies were applied to generate other constructs. For point mutation, a site-directed mutagenesis kit (New England Biolabs Q5) was used. When making fusions between the different linker domains and POU_H_, we maintained the “RKRKR” sequence at the N-terminus of Oct4 POU_H_, which has been shown to be important for reprogramming (Jerabek et al. 2017).

### iPSC generation

293T cells were transfected with mSTEMCCA encoding FLAG-tagged Oct4, Oct1 and different domain-swaps and mutants as well as the packaging plasmids (pMDLg/pRRE and pRSV-Rev) and envelope plasmid (VSV-G) using polyethylenimine. Lentiviral supernatants were filtered, 8 μg/mL polybrene (Sigma) was added, and the mixture used to transduce passage 3 primary MEFs expressing EGFP from the endogenous *Pou5f1* locus. The MEFs were prepared from E12.5 mouse embryos (Jackson Labs strain #008214, B6;129S4-Pou5f1^tm2Jae^/J) and were seeded in 6-well plates the night before transduction at a density of 1×10^5^ cells/well. Cells were cultured in ESC medium (above). GFP^+^ iPSC colony emergence was quantified after 12 days using an Olympus IX-71 epifluorescence microscope. Lentivirus titers were established using two techniques. For the experiment shown in Supplemental Fig. S2B, a kit measuring p24 was used (QuickTiter Lentivirus Titer Kit, Cell Biolabs). For the experiment shown in Supplemental Fig. S4C, a kit measuring conserved areas of viral RNA was used (Lenti-X qRT-PCR, Takara Bio).

### Immunoblotting and immunofluorescence

Antibodies for immunoblotting were: Oct4, α-tubulin and β-actin, Santa Cruz; Oct1, Bethyl; Nanog and Sox2, GeneTex; GAPDH, EMD Millipore; FK2 (pan-Ub), Enzo Life Sciences; Ub-K48, Cell Signaling. All immunoblots are shown in uncropped form in Supplemental Fig. S14-17. For immunofluorescence, testes from both *Oct4*^Jky/Jky^ mouse and an *Oct4*^+/+^ littermate control were weighed and fixed using 4% formaldehyde. The paraffinized testes were sectioned into 5 μm slices. Antibodies for immunofluorescence were: mouse anti-SYCP3, Santa Cruz; goat anti-LIN28A, R&D Systems; rabbit anti-phospho-H2AX, Cell Signaling; Alex Fluor 647 donkey anti-mouse IgG, Alex Fluor 488 donkey anti-rabbit IgG and Alex Fluor 594 donkey anti-goat IgG, ThermoFisher Scientific. Data were imaged using a Nikon ECLIPSE Ti inverted microscope with Nikon Elements AR software.

### Electrophoretic mobility shift assay

EMSA was performed using published methods (Kang et al. 2009; Ferraris et al. 2011). Diamide (Sigma) was added at final concentration of up to 300 μM on ice during reaction assembly. 5’Cy5-labeled double-stranded DNA probes were as follows: wild type octamer, 5’-TGTCGAATGCAAATCACT-3’; mutant octamer, 5’-TGTCGAATGCAAGCCACT-3’. Reactions were incubated for 30 min at room temperature prior to gel loading. For experiments involving reversal of oxidative inhibition, thioredoxin (Sigma) was supplied at a final concentration of 0 to 60 μM after 15 min incubation with diamide, and both thioredoxin-treated and control samples were incubated for a further 15 min at 37°C prior to gel loading. DNA was visualized using a Molecular Dynamics Typhoon system.

### Recombinant Oct4 protein

Wild type and C48S Oct4 constructs were generated bearing dual C-terminal FLAG and Twin-Strep tags. A g-block (IDT) fragment coding for the Twin-Strep and FLAG epitope tags was inserted between the *Nco*I and *Kpn*I sites in the transient expression vector pACE-MAM2 (Geneva Biotech). The resultant construct was then digested with *Nco*I and *Pvu*II to insert the g-block fragment coding for either wild type or C48S murine Oct4. After Sanger sequencing to confirm veracity, the plasmids were transfected into Expi293F cells (ThermoFisher) using ExpiFectamine 293 Tranfection Kit (ThermoFisher) following the manufacturer instructions. After 2 days, transfected cells were harvested and washed with ice-cold PBS. Cells were resuspended in Buffer A (20 mM HEPES pH 8.0, 1.5 mM MgCl_2_, 10 mM KCl, 0.25% NP-40, 0.5 mM DTT, 2x protease inhibitor cocktail/Roche), incubated for 10 min on ice, and homogenized using a Dounce homogenizer (Wheaton). The homogenate was centrifuged at 4°C for 5 min at 3,800×*g*). Pelleted nuclei were resuspended in Buffer C (20 mM HEPES pH 8.0, 25% glycerol, 1.5 mM MgCl_2_, 420 mM KCl, 0.25% NP-40, 0.2 mM EDTA, 0.5 mM DTT, 2x protease inhibitor cocktail) and Dounce homogenized again. Nuclei were extracted for 30 min at 4°C with end-to-end rotation, then centrifuged at 4°C for 30 min at 20,000×*g*. The nuclear extract was then subjected to immunoprecipitation with anti-FLAG (M2)-agarose beads (Sigma). Bead-bound proteins were eluted with 200 ng/µL 3×FLAG peptide (Sigma) by incubating at 4°C for 30 min. The protein was pure and monodisperse as verified using Coomassie blue staining and size exclusion chromatography.

### Generation of Oct4^C48S^ mouse ESCs

Mutant ESCs with a single-base missense mutation that converts Cys48 to Ser were generated using templated CRISPR. A modified synthetic sgRNA (Synthego) was designed to target Cas9 and cutting near the target nucleotide, 5’ GGCCTCGAAGCGA<u>C</u>AGATGG 3’, where the underlined C indicates the target nucleotide. The sgRNA was complexed with Cas9 protein (IDT) to form a ribonucleoprotein (RNP) complex that was electroporated with program CG-104 into 6×10^4^ mouse ESCs using a LONZA 4D-Nucleofector X-unit. The RNP was co-electroporated with a corresponding single-stranded oligodeoxynucleotide (ssODN) with 5’ and 3’ phosphorothioate modifications, 5’-ctcactctgtttgatcggcctttcagGAAAGGTGTTCAGCCAGACCA**CCATCT<u>C</u>T<u>A</u>G<u>A</u>TTCGAGGCC**TTGCAGCTCAGCCTTAAGAACATGTGTAAGCTG-3’ where lower-case denotes intronic sequence and the sgRNA site is in bold. The ssODN contained the target mutation (underlined), silent mutations in the seed region (double underlined) to block CRISPR re-cutting after incorporation of the donor and to generate a unique *Xba*I restriction enzyme site for screening purposes. Incorporation of the ssODN sequence was identified via PCR and restriction enzyme digestion and confirmed by sequencing of the 709 bp region flanking the mutation. Mutant clones were generated from a pure C57BL/6 ESC line, a 129/BL6 hybrid line expressing constitutive GFP and a reporter line used to de-differentiate ESCs into a 2C-like state. Clones were tested by karyotype and any clones with <70% normal diploid content were excluded.

### RNA-seq

ESCs were first depleted by feeder cells by incubation on plates coated with gelatin. Parental and derived Oct4^C48S^ ESC lines were differentiated in media lacking LIF and containing 1.5mg/L RA for 2 days. Undifferentiated and RA-differentiated cells were collected, and RNA was prepared using a kit (RNeasy, Qiagen). RNA poly-A selected for mRNA using NEBNext rRNA Depletion Kit v2 (Human, Mouse, Rat) (E7400). Bulk RNA was processed and sequenced as reported previously (Perovanovic et al. 2023). Briefly, RNASeq reads were aligned to GRCh38/hg38 using STAR version 2.7.9a. Read counts were normalized using the DESeq2 analysis package. The DESeq2-regularized log transformation was applied to counts, which preferentially shrinks the overall variance among low abundant transcripts. The high-throughput data are available on the gene expression omnibus (GEO) server under series record GSE241332. The data are accessible for download through private token gtyfyseqljufjwz while in private status. Differential genes were identified using 2.5-fold change and *p*<0.001 as cutoffs. GO terms were identified using PANTHER (Mi et al. 2013).

### Teratoma formation

Feeder-depleted parental and derived Oct4^C48S^ ESC lines were injected subcutaneously into contralateral flanks of 6-8 week-old female NOD.*Rag1^-/-^Il2rg^-/-^* (NRG, Jackson labs) mice to generate teratomas. 1×10^6^ cells were implanted. After two weeks, mice were humanely euthanized, and tumors were excised. After fixation, embedding and sectioning, slides were stained with H&E and evaluated in a blinded fashion.

### Chimeric mouse assays

Targeted or parental ESCs were collected and resuspended in injection medium consisting of DMEM with HEPES (ThermoFisher) supplemented with 10% FBS (Peak Serum) and 0.1 mM 2-mercaptoethanol (Sigma). C57B6 clones were injected into albino B6 blastocysts, while the G4N and C4 clones (B6:129 and 129 in origin, respectively) were injected into C57B6 blastocysts. E3.5 blastocysts were used. 10-25 ESCs were injected into each blastocyst using micromanipulators (Eppendorf) and a Leica inverted microscope. After culturing injected blastocysts in KSOM (CytoSpring), approximately 12 to 18 blastocysts were surgically transferred into the uterine horns of day 2.5 pseudopregnant mice or the oviduct of day 0.5 pseudopregnant mice. Mice were scored based on coat and eye color by members of the transgenic core facility at weaning.

### Janky mice

Mutant C57BL6/J mice with a single-base missense mutation that converts Cys48 to Ser (*Jky* mice) were generated using the same templated CRISPR strategy used to generate mutant ESCs. CRISPR RNPs were introduced into single cell C57BL6/J mouse embryos using pronuclear injection. Founder mice that carried the targeted mutation were backcrossed to WT and the N1 generation was sequenced to confirm the presence of the heterozygous allele.

## Supporting information

Supplemental Move S1

Supplemental Move S2

Supplemental Move S3

Supplemental Move S4

Supplemental Move S5

Supplemental Move S6

## Acknowledgements

We thank S. Buckley for critical reading of the manuscript. We thank C. Davey, H. Lan and the University of Utah Mouse Transgenic and Gene Targeting Core as well as the Mutation Generation and Detection Core for assistance with CRISPR-edited ESC lines, chimerism assays, karyotyping and the generation of germline mutant mice. We thank B. Dalley and the High-Throughput Genomics Core facility, and Q. Li and the Bioinformatics Analysis core facility for assistance with RNA-seq. We thank D. Lum and the Preclinical Research Resource core facility for assistance with teratoma formation. We thank J. Marvin and the Flow Cytometry Core, and X. Wang and the Cell Imaging Core. We thank B. Weaver for the gift of the C4 ESC line. This work was supported by NIH/NIGMS grant (R01GM122778) to DT.

## Author contributions

D.T. conceived the study. Z.S., Y.W. and D.T. designed experiments. Z.S., Y.W. and M.B.C. generated new reagents for the study. Z.S., Y.W., A.M., C.Y., B.R.C. and K.J.E. performed experiments and/or analyzed data. K.J.E. performed blinded pathological analysis of teratoma samples. Z.S., Y.W. and A.M. generated figures. All authors contributed to the writing of the manuscript and read and approved the final version of the manuscript prior to submission for publication.

## Supplemental figure legends

**Figure S1.** Effect of N-terminal FLAG-tag on iPSC formation. (*A*) Similar to Fig. 1C, except cells were transduced with lentiviral vectors expressing untagged and N-terminal rather than C-terminal FLAG-tagged Oct4. (*B*) Anti-FLAG and Oct4 immunoblots showing expression of tagged and untagged Oct4. α-tubulin is shown as a loading control.

**Figure S2.** GFP^+^ iPSC numbers over time using different Oct1-Oct4 chimeric constructs. (*A*) Similar to Fig. 1D except multiple time points are shown. Wild type Oct4 is shown as a positive control. Results show an average of experiments performed in biological triplicate. Error bars show ±standard deviation. (*B*) Titers for six of the viral supernatants used in Fig.1 and in (A) were tested for p24 levels using a kit. Averages of N=2 independent experiments are shown. TU=transduction units.

**Figure S3.** iPSC colony morphology and GFP intensity using different chimeric and mutant reprogramming constructs. (*A*) Phase-contrast and epifluorescence images are shown of example iPSC colonies generated using the constructs shown in Fig. 2. Scale bar=200 μm. (B) For each construct in (A), GFP intensity from N=X colonies from three different wells was quantified by dividing the total GFP pixel intensity by the area of each colony, generating an average GFP intensity per unit area (arbitrary units). Values for each well were then averaged. Error bars denote ±standard deviation.

**Figure S4.** The effect of Cys48 on iPSC colony formation is not dependent on kinetics or on using Oct4-GFP MEFs. (*A*) GFP^+^ iPSC numbers over time using different Oct1-Oct4 chimeric constructs. Similar to Fig. 2A except multiple time points are shown. Wild type Oct4 is shown as a positive control. Results show an average of experiments performed in biological triplicate. Error bars show ±standard deviation. (*B*) An iPSC colony formation assay using non-GFP primary MEFs and alkaline phosphatase staining rather than GFP positivity and pluripotent morphology to score iPSC formation. N=3. Error bars show ±standard deviation. (*C*) Titers of the viral supernatants used in (B) were measured based on viral RNA levels using a kit. Averages of N=3 replicates are shown. Error bars show ±standard deviation. IFU=infectious units.

**Figure S5.** Oxidative inhibition of Oct4 DNA binding can be reversed with the addition of thioredoxin. EMSA is shown similar to Fig. 3C, except that after 15 min incubation with diamide, thioredoxin was added at the indicated concentrations. Reactions were incubated for a further 15 min at 37°C prior to native electrophoresis.

**Figure S6.** Purification of recombinant wild type and C48S mutant Oct4 from expi-293T cells. Oct4 was FLAG-tagged at the C-terminus. Protein was purified using beads covalently coupled to anti-FLAG antibodies. Lysates were prepared 48 hr post-transfection.

**Figure S7.** H_2_O_2_ and MG-132 treatment of primary MEFs largely maintains viability. Phase-contrast images are shown of Oct4-GFP MEFs treated with the same concentrations and timepoints of H_2_O_2_ and MG-132 used in Fig. 3E. Scale bar=200 μm.

**Figure S8**. Oct4 C48S promotes Oct4 ubiquitylation. Similar to Fig. 3F except additional antibodies were used for immunoblotting: an antibody detecting K63-linked Ub chains (Cell Signaling) and a pan-Oct4 antibody as a positive control (Enzo Life Sciences). Oct4 was Twin-Strep-and FLAG-tagged at the C-terminus.

**Figure S9.** The effects of Oct4 C48S mutation are not parent line- or clone-specific. (*A*) High-resolution melting analysis of parental, homozygous mutant and Indel clones generated by CRISPR targeting of *Pou5f1* in mouse ESCs. The majority of clones were Indels and a smaller number of clones were homozygous mutant. Heterozygous and unedited (wild-type) lines were not recovered by the procedure. (*B*) Similar to Fig. 4D except using alkaline phosphatase staining. (C) Similar to Fig. 4B except a different clone from the same parent line (#2) was tested. (*C*) Similar to Fig. 4F except a different clone from the same parent line (#2) was tested. (*D*) Quantification from N=3 immunoblots. Background was subtracted from individual bands, and intensity for each protein was divided by H3 loading control to generate a measure of relative protein level in arbitrary units. Error bars denote ±standard deviation. Quantification was performed with Image J software. (*E*) Similar to Fig. 4F except the same clone was tested against parent ESCs that had been simultaneously electroporated without CRISPR RNP (“zap” control). (*F*) Similar to Fig. 4F except a different clone from the same parent line (#2) was tested. (*G*) Similar to Fig. 4F except a different clone (#18) from a different parent line (G4-N) was tested.

**Figure S10.** RNA-seq analysis of differentiating parental and Oct4^C48S^ ESCs. (*A*) Similarity between bulk RNA-seq replicates and conditions. Clustering of the RNA-seq replicates using Euclidean distance is shown. (*B*) The union of the top 100 most differentially expressed genes upon differentiation for both genotypes (150 genes total) was subjected to hierarchical clustering and displayed as a heat map. Example genes are shown at right.

**Figure S11.** Additional G4-N clone #18 teratoma images similar to Fig.6D. Asterisk represents necrotic regions.

**Figure S12.** The decreased differentiation of teratomas from Oct4^C48S^ ESCs normalizes in larger tumors at later timepoints. (*A*) Representative H&E-stained sections of tumors dissected from parental and mutant teratomas at day 28. Regions surrounded by black lines highlight immature neuroepithelial elements. Blue lines highlight mature epithelium (squamous, respiratory, and/or gastroinstestinal). Red asterisk represents mature neural regions. Green lines show mature cartilage. Yellow lines show mature fat. Orange lines show yolk sac (immature). (*B*) Average percent immature component for day 28 parental homozygous mutant teratomas. N=3 individual tumors. Error bars denote ±standard deviation. (*C*) Similar analysis performed for a different parent and homozygous Oct4^C48S^ mutant BL/6 ESC pair grown as teratomas for 28 days. N=5.

**Figure S13.** Abnormal testes in *Jky* homozygous mice. (*A*) Gross images of dissected testes from a 2.5 month-old male *Jky*/*Jky* mouse and +/+ littermate control. Averaged weights of two testes from each animal are also shown. (*B*) H&E staining of FFPE testes sections from *Jky*/*Jky* and +/+ littermate control testes. (*C*) Representative immunofluorescence images showing abnormal spermatogenesis and germ cell loss in testis sections from the *Oct4*^Jky/Jky^ mouse: gamma H2AX (green); LIN28A (Red); SYCP3 (violet); and DNA (Cyan). Box #1 highlights examples of superficially normal spermatogenesis, while #2 highlights example abnormal localization of spermatocytes and germ cell loss. Arrow points to abnormal localization of spermatocytes within interstitial regions.

**Figure S14.** Uncropped immunoblots for Figs. 1-3.

**Figure S15.** Uncropped immunoblots for Fig. 4.

**Figure S16.** Uncropped immunoblots for Suppl. Fig. S1 and S8.

**Figure S17.** Uncropped immunoblots for Suppl. Fig. S9.

**Movie S1.** Example of beating cells derived from parent EBs plated and differentiated into cardiomyocytes.

**Movie S2.** Example of beating cells derived from parent EBs plated and differentiated into cardiomyocytes.

**Movie S3.** Example of non-beating cells derived from mutant #2 EBs plated and differentiated into cardiomyocytes.

**Movie S4.** Example of non-beating cells derived from mutant #2 EBs plated and differentiated into cardiomyocytes.

**Movie S5.** Example of non-beating cells derived from mutant #46 EBs plated and differentiated into cardiomyocytes.

**Movie S6.** Example of non-beating cells derived from mutant #46 EBs plated and differentiated into cardiomyocytes.

