## Supplemental Figures S1-S17 for "Oct4 redox sensitivity potentiates reprogramming and differentiation"

[1]Department of Pathology,

Supplemental material includes:
Supplemental Figs. S1-S17 (.pdf)
Supplemental Tables S1-S3 (.xlsx)
Supplemental Movies S1-S6 (.mp4)

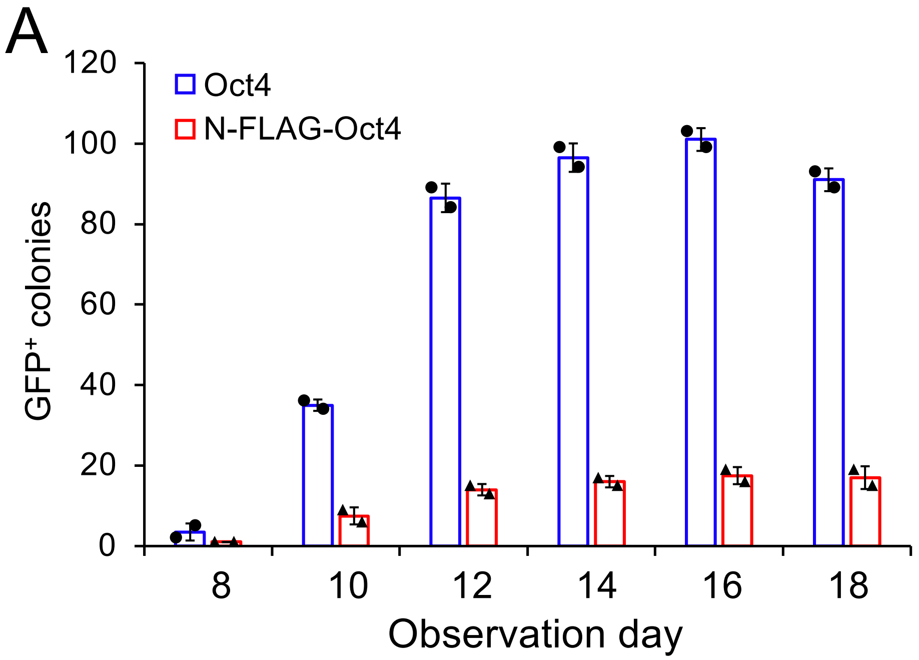
A
120
100
80
60
40
20
0
Oct4
N-FLAG-Oct4
GFP+ colonies
8
10
12
14
16
18
Observation day

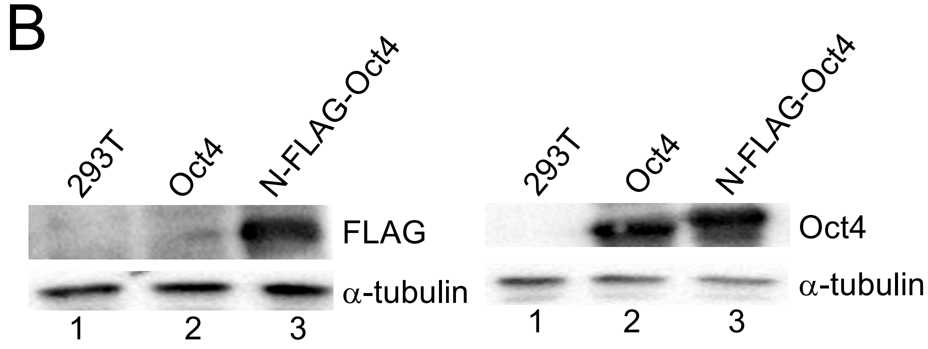
B
293T
Oct4
N-FLAG-Oct4
FLAG
α-tubulin
1
2
3
293T
Oct4
N-FLAG-Oct4
Oct4
α-tubulin
1
2
3

**Figure S1.** Effect of N-terminal FLAG-tag on iPSC formation. (*A*) Similar to Fig. 1C, except cells were transduced with lentiviral vectors expressing untagged and N-terminal rather than C-terminal FLAG-tagged Oct4. (*B*) Anti-FLAG and Oct4 immunoblots showing expression of tagged and untagged Oct4. α-tubulin is shown as a loading control.

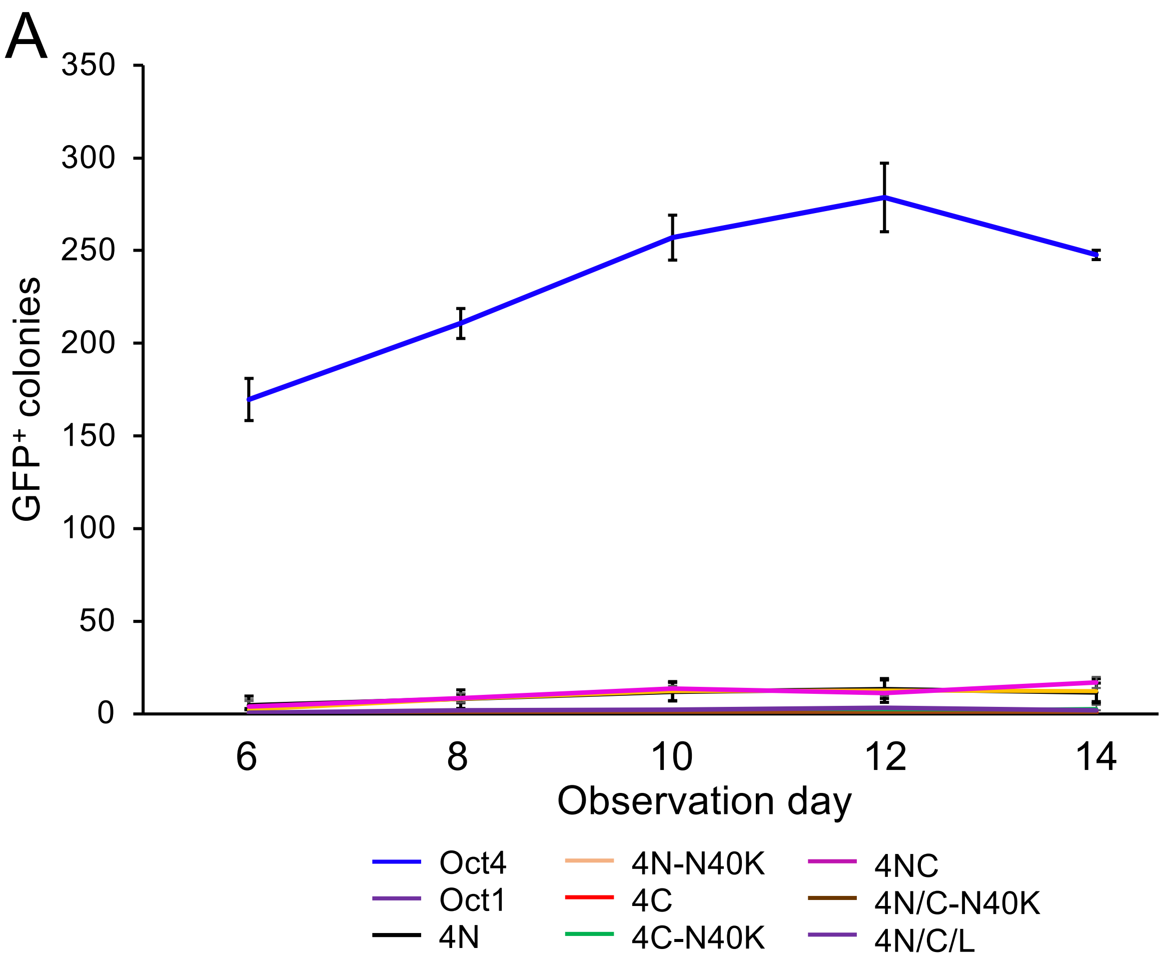
A
GFP⁺ colonies
350
300
250
200
150
100
50
0
6
8
10
12
14
Observation day
Oct4
Oct1
4N
4N-N40K
4C
4C-N40K
4NC
4N/C-N40K
4N/C/L

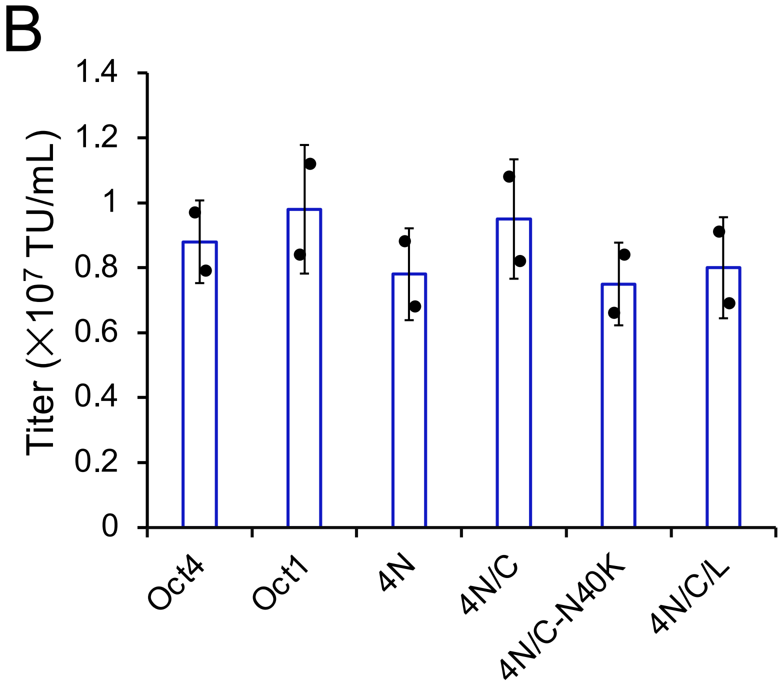
B
Titer (×10⁷ TU/mL)
1.4
1.2
1
0.8
0.6
0.4
0.2
0
Oct4
Oct1
4N
4N/C
4N/C-N40K
4N/C/L

**Figure S2.** $GFP^+$ iPSC numbers over time using different Oct1-Oct4 chimeric constructs. (*A*) Similar to Fig. 1D except multiple time points are shown. Wild type Oct4 is shown as a positive control. Results show an average of experiments performed in biological triplicate. Error bars show ±standard deviation. (*B*) Titers for six of the viral supernatants used in Fig.1 and in (A) were tested for p24 levels using a kit. Averages of N=2 independent experiments are shown. TU=transduction units.

Shen et al. Figure S3

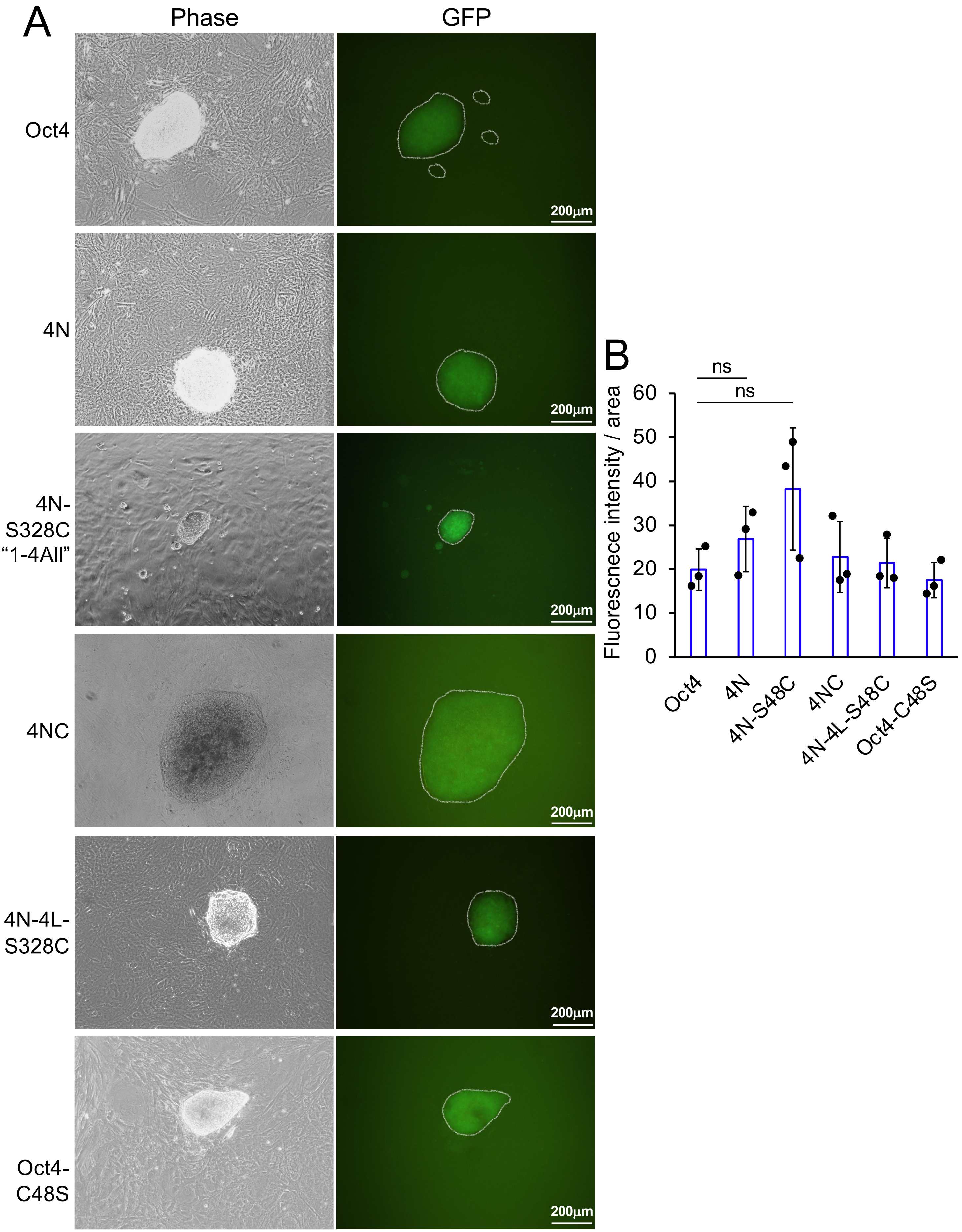

**Figure S3.** iPSC colony morphology and GFP intensity using different chimeric and mutant reprogramming constructs. (*A*) Phase-contrast and epifluorescence images are shown of example iPSC colonies generated using the constructs shown in Fig. 2. Scale bar=200 μm. (*B*) For each construct in (A), GFP intensity from N=3 colonies from three different wells was quantified by dividing the total GFP pixel intensity by the area of each colony, generating an average GFP intensity per unit area (arbitrary units). Values for each well were then averaged. Error bars denote ±standard deviation.

Shen et al. Figure S4

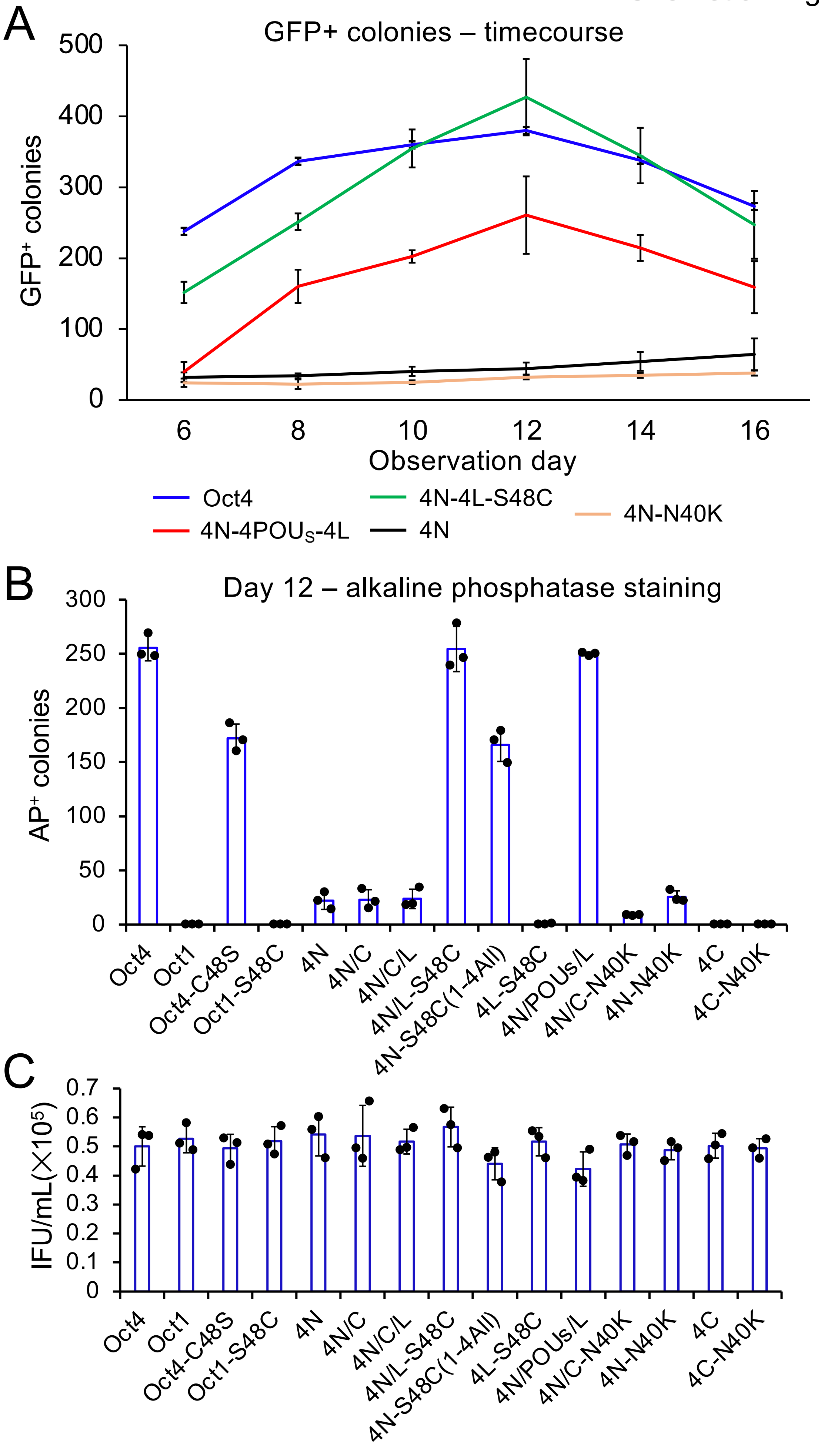

**Figure S4.** The effect of Cys48 on iPSC colony formation is not dependent on kinetics or on using Oct4-GFP MEFs. (*A*) $GFP^+$ iPSC numbers over time using different Oct1-Oct4 chimeric constructs. Similar to Fig. 2A except multiple time points are shown. Wild type Oct4 is shown as a positive control. Results show an average of experiments performed in biological triplicate. Error bars show ±standard deviation. (*B*) An iPSC colony formation assay using non-GFP primary MEFs and alkaline phosphatase staining rather than GFP positivity and pluripotent morphology to score iPSC formation. N=3. Error bars show ±standard deviation. (*C*) Titers of the viral supernatants used in (B) were measured based on viral RNA levels using a kit. Averages of N=3 replicates are shown. Error bars show ±standard deviation. IFU=infectious units.

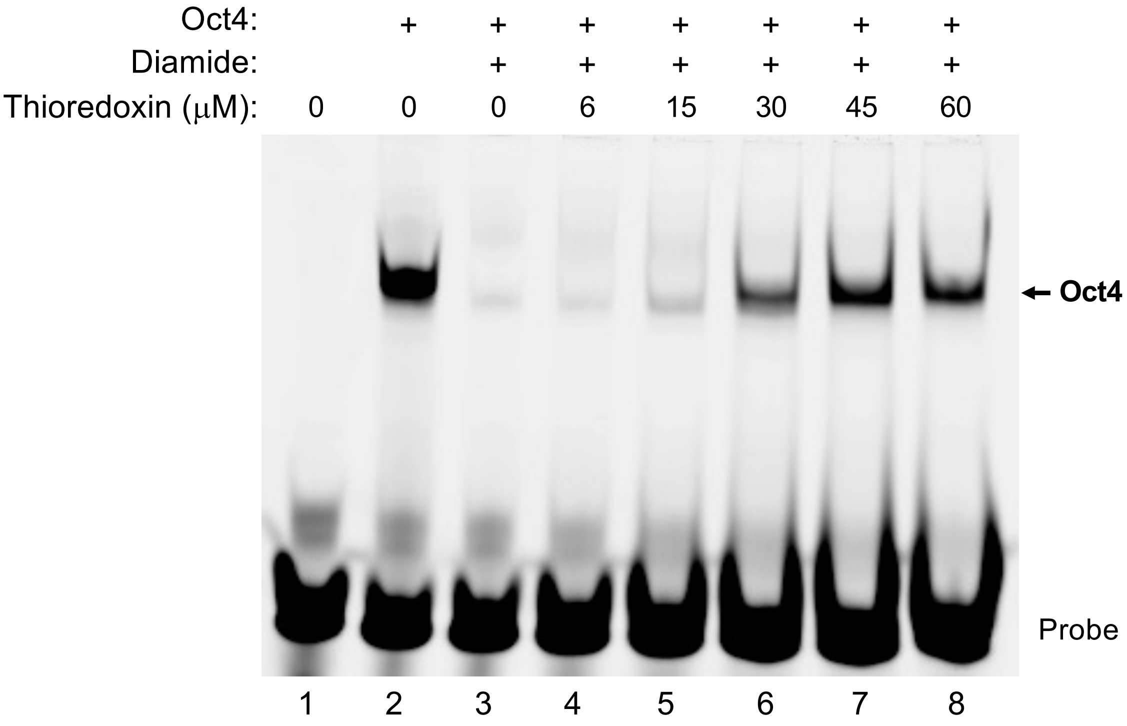
Oct4: + + + + + + +
Diamide: + + + + + +
Thioredoxin (μM): 0 0 0 6 15 30 45 60
← Oct4
Probe
1 2 3 4 5 6 7 8

**Figure S5.** Oxidative inhibition of Oct4 DNA binding can be reversed with the addition of thioredoxin. EMSA is shown similar to Fig. 3C, except that after 15 min incubation with diamide, thioredoxin was added at the indicated concentrations. Reactions were incubated for a further 15 min at 37°C prior to native electrophoresis.

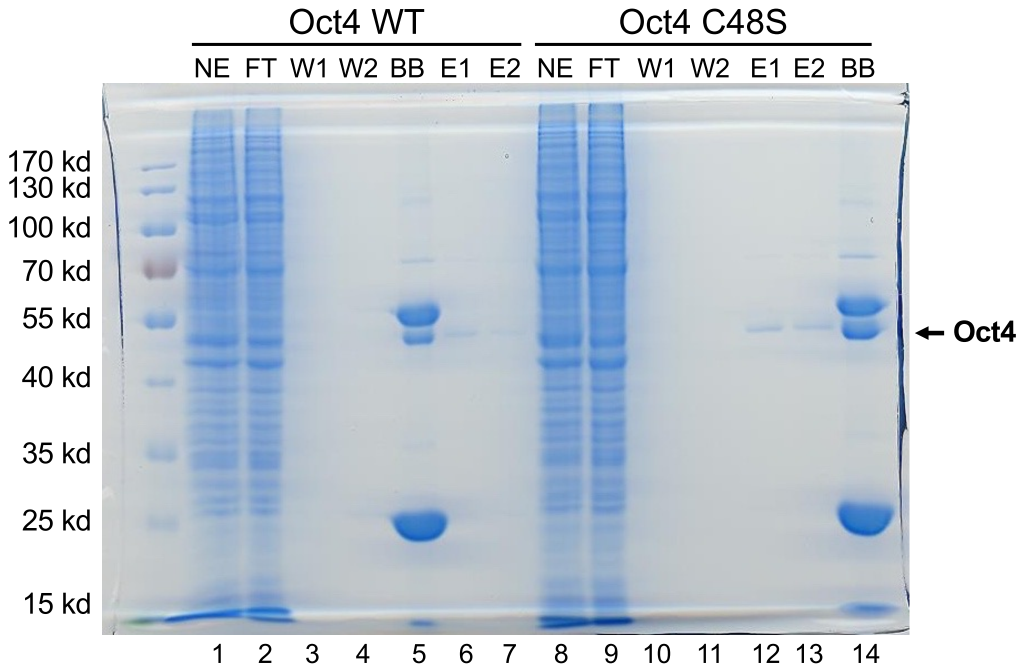


| | |
|---|---|
| NE | Nuclear extract |
| FT | Flow through |
| W | Wash |
| BB | Boiled beads |
| E | Elution |

**Figure S6.** Purification of recombinant wild type and C48S mutant Oct4 from expi-293T cells. Oct4 was FLAG-tagged at the C-terminus. Protein was purified using beads covalently coupled to anti-FLAG antibodies. Lysates were prepared 48 hr post-transfection.

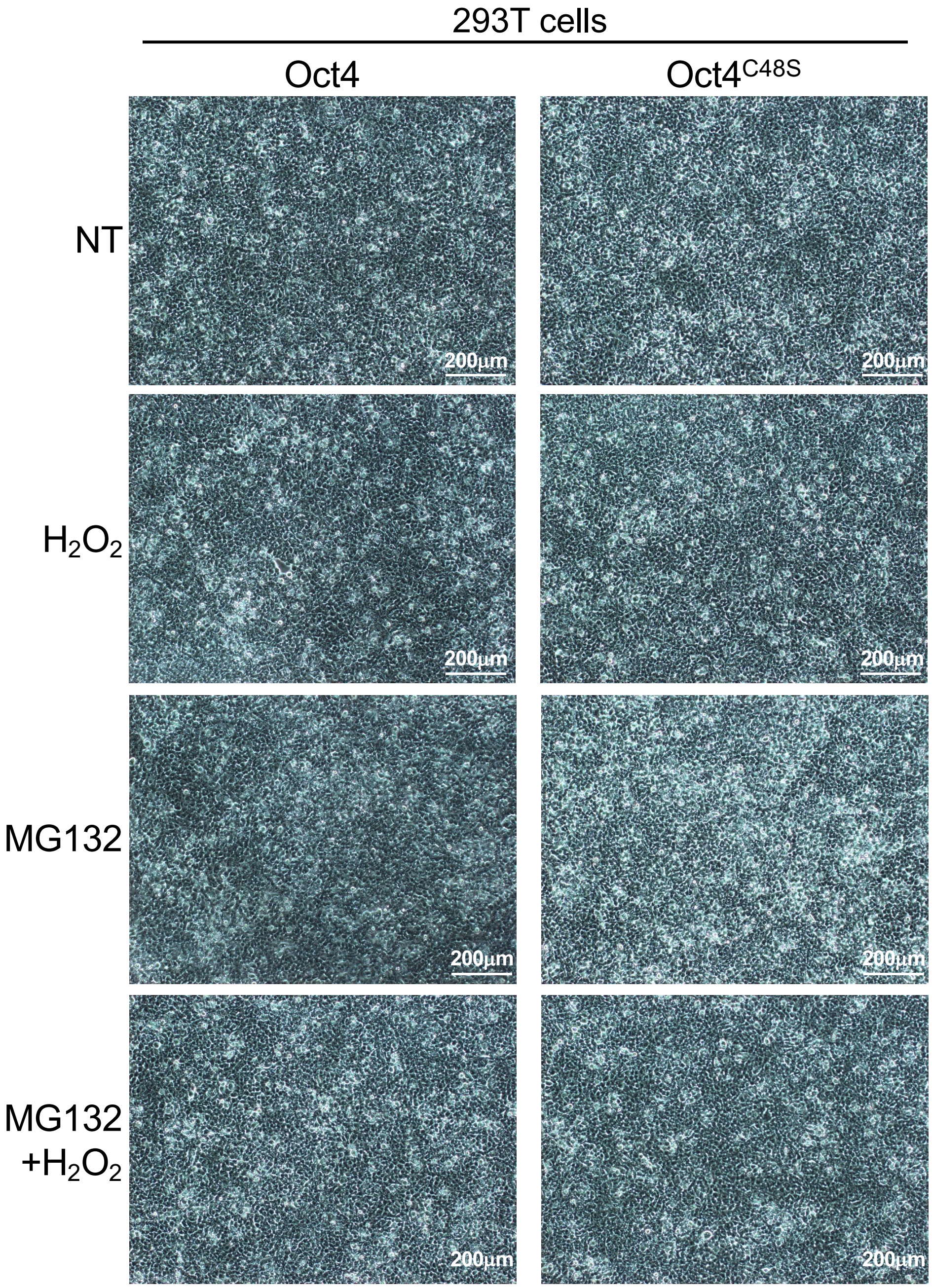
293T cells
Oct4
Oct4C48S
NT
H2O2
MG132
MG132 +H2O2
200μm

**Figure S7.** $H_2O_2$ and MG-132 treatment of primary MEFs largely maintains viability. Phase-contrast images are shown of Oct4-GFP MEFs treated with the same concentrations and timepoints of $H_2O_2$ and MG-132 used in Fig. 3E. Scale bar=200 μm.

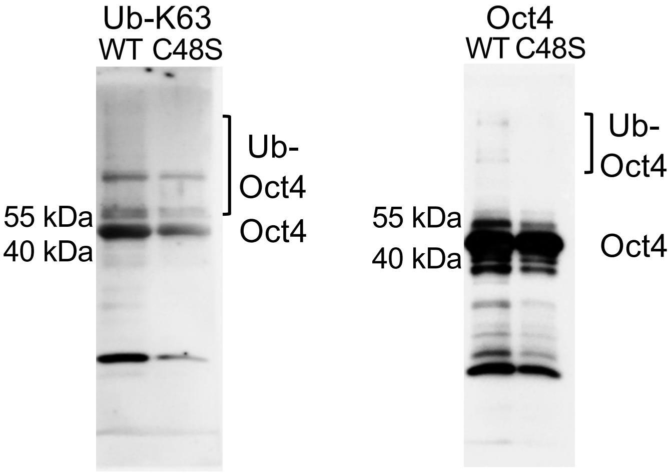
Ub-K63
WT C48S
Ub-
Oct4
55 kDa
40 kDa
Oct4
Oct4
WT C48S
Ub-
Oct4
55 kDa
40 kDa
Oct4

**Figure S8.** Oct4 C48S promotes Oct4 ubiquitylation. Similar to Fig. 3F except additional antibodies were used for immunoblotting: an antibody detecting K63-linked Ub chains (Cell Signaling) and a pan-Oct4 antibody as a positive control (Enzo Life Sciences). Oct4 was Twin-Strep- and FLAG-tagged at the C-terminus.

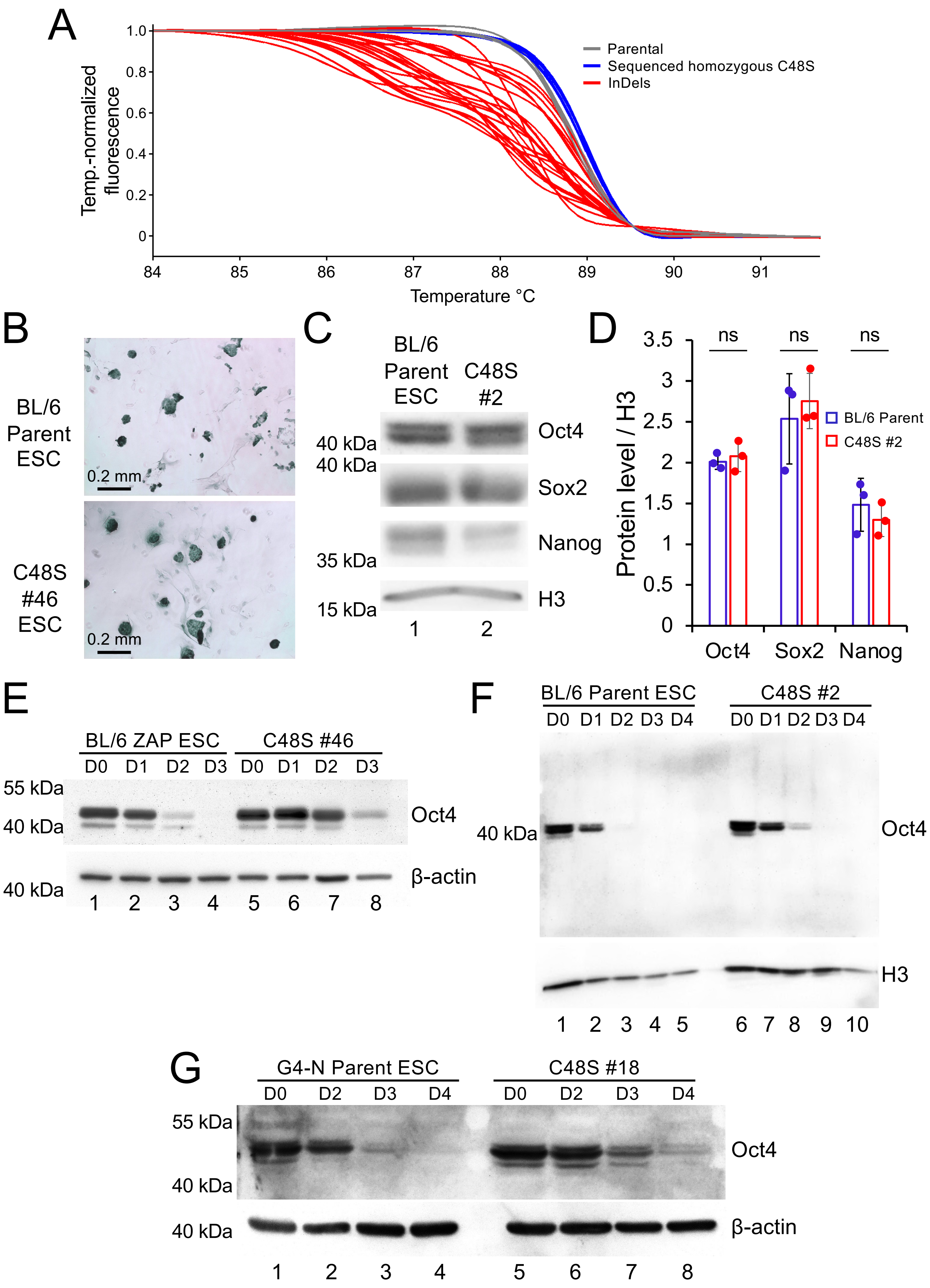
A
Temp.-normalized fluorescence
1.0
0.8
0.6
0.4
0.2
0
84
85
86
87
88
89
90
91
Temperature °C
Parental
Sequenced homozygous C48S
InDels
B
BL/6 Parent ESC
C48S #46 ESC
0.2 mm
0.2 mm
C
BL/6 Parent ESC
C48S #2
40 kDa
40 kDa
35 kDa
15 kDa
Oct4
Sox2
Nanog
H3
1
2
D
Protein level / H3
3.5
3
2.5
2
1.5
1
0.5
0
ns
ns
ns
BL/6 Parent
C48S #2
Oct4
Sox2
Nanog
E
BL/6 ZAP ESC
C48S #46
D0 D1 D2 D3 D0 D1 D2 D3
55 kDa
40 kDa
40 kDa
Oct4
β-actin
1 2 3 4 5 6 7 8
F
BL/6 Parent ESC
C48S #2
D0 D1 D2 D3 D4 D0 D1 D2 D3 D4
40 kDa
Oct4
H3
1 2 3 4 5 6 7 8 9 10
G
G4-N Parent ESC
C48S #18
D0 D2 D3 D4 D0 D2 D3 D4
55 kDa
40 kDa
40 kDa
Oct4
β-actin
1 2 3 4 5 6 7 8

**Figure S9.** The effects of Oct4 C48S mutation are not parent line- or clone-specific. (*A*) High-resolution melting analysis of parental, homozygous mutant and Indel clones generated by CRISPR targeting of *Pou5f1* in mouse ESCs. The majority of clones were Indels and a smaller number of clones were homozygous mutant. Heterozygous and unedited (wild-type) lines were not recovered by the procedure. (*B*) Similar to Fig. 4D except using alkaline phosphatase staining. (*C*) Similar to Fig. 4B except a different clone from the same parent line (#2) was tested. (*D*) Quantification from N=3 immunoblots. Background was subtracted from individual bands, and intensity for each protein was divided by H3 loading control to generate a measure of relative protein level in arbitrary units. Error bars denote ±standard deviation. Quantification was performed with Image J software. (*E*) Similar to Fig. 4F except the same clone was tested against parent ESCs that had been simultaneously electroporated without CRISPR RNP ("zap" control). (*F*) Similar to Fig. 4F except a different clone from the same parent line (#2) was tested. (*G*) Similar to Fig. 4F except a different clone (#18) from a different parent line (G4-N) was tested.

A

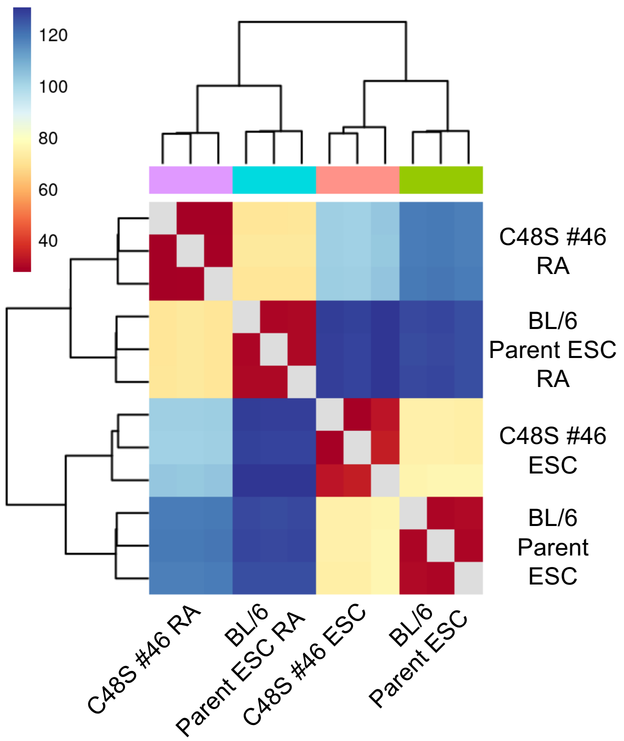
120
100
80
60
40
C48S #46 RA
BL/6 Parent ESC RA
C48S #46 ESC
BL/6 Parent ESC
C48S #46 RA
BL/6 Parent ESC RA
C48S #46 ESC
BL/6 Parent ESC

B

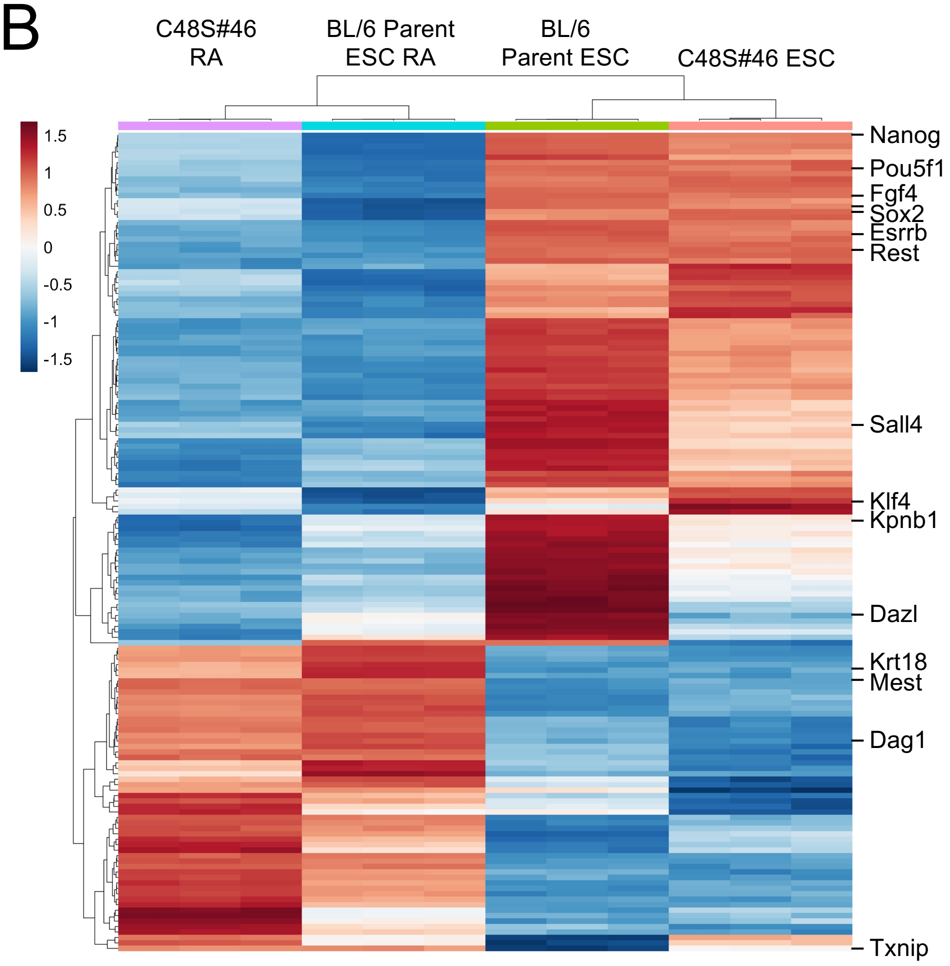
C48S#46 RA
BL/6 Parent ESC RA
BL/6 Parent ESC
C48S#46 ESC
1.5
1
0.5
0
-0.5
-1
-1.5
Nanog
Pou5f1
Fgf4
Sox2
Esrrb
Rest
Sall4
Klf4
Kpnb1
Dazl
Krt18
Mest
Dag1
Txnip

**Figure S10.** RNA-seq analysis of differentiating parental and Oct4$^{C48S}$ ESCs. (*A*) Similarity between bulk RNA-seq replicates and conditions. Clustering of the RNA-seq replicates using Euclidean distance is shown. (*B*) The union of the top 100 most differentially expressed genes upon differentiation for both genotypes (150 genes total) was subjected to hierarchical clustering and displayed as a heat map. Example genes are shown at right.

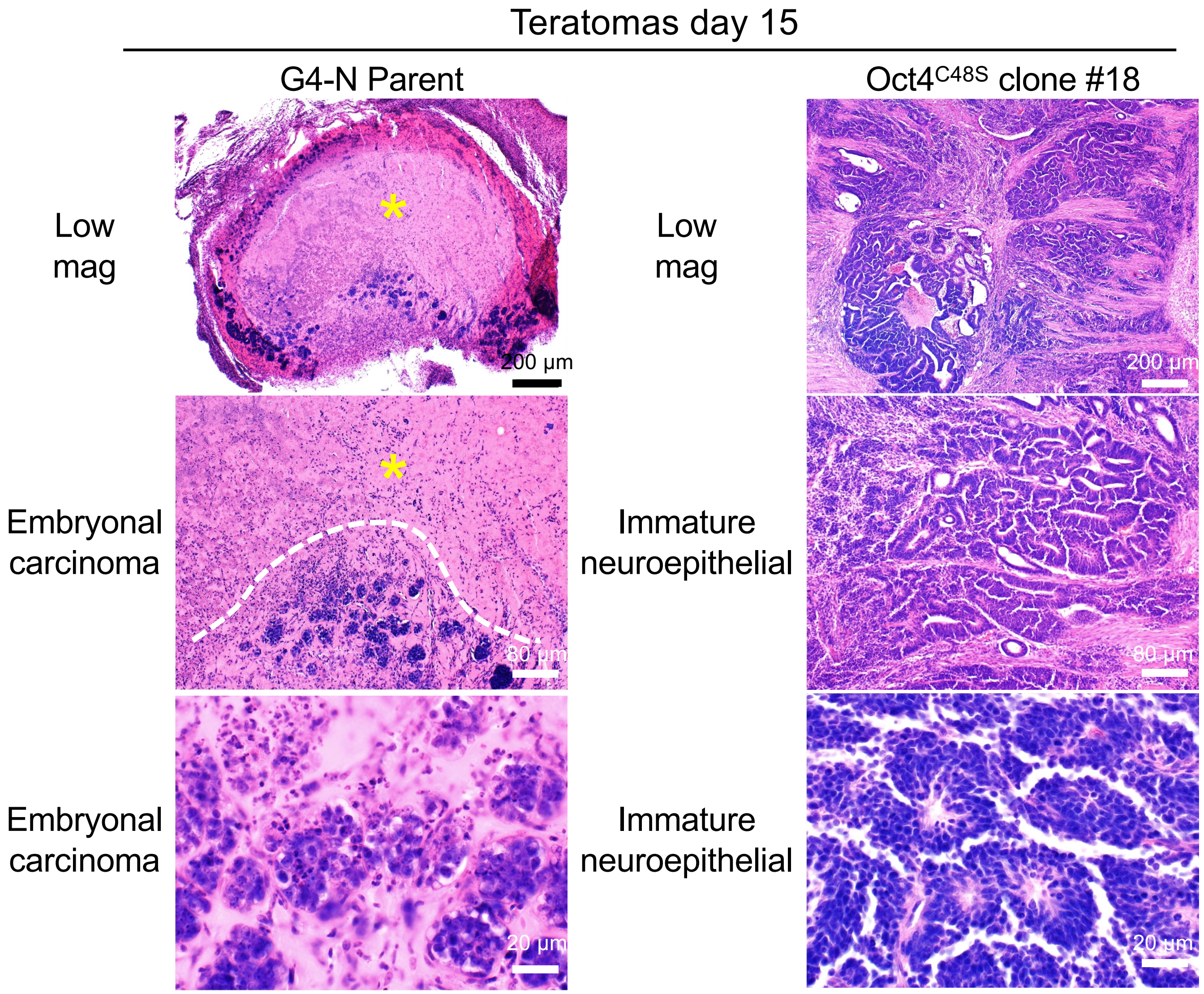
Teratomas day 15
G4-N Parent
$Oct4^{C48S}$ clone #18
Low mag
Low mag
200 µm
200 µm
Embryonal carcinoma
Immature neuroepithelial
80 µm
80 µm
Embryonal carcinoma
Immature neuroepithelial
20 µm
20 µm

**Figure S11.** Additional G4-N clone #18 teratoma images similar to Fig.6D. Asterisks represent necrotic regions.

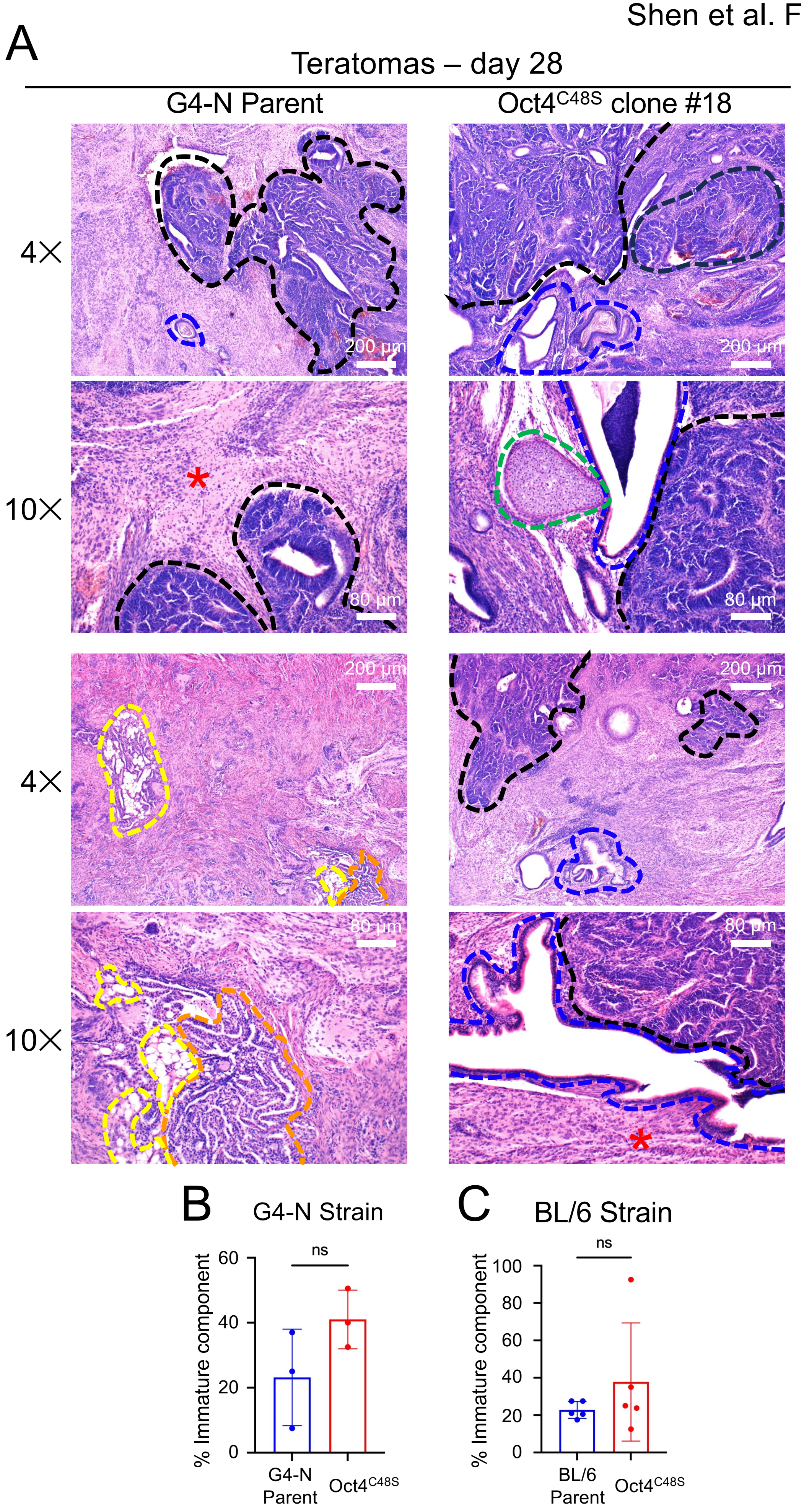
A
Teratomas – day 28
G4-N Parent
Oct4C48S clone #18
4×
10×
4×
10×
200 µm
80 µm
200 µm
80 µm
200 µm
200 µm
80 µm
80 µm
B
G4-N Strain
ns
% Immature component
60
40
20
0
G4-N Parent
Oct4C48S
C
BL/6 Strain
ns
% Immature component
100
80
60
40
20
0
BL/6 Parent
Oct4C48S

**Figure S12.** The decreased differentiation of teratomas from Oct4$^{C48S}$ ESCs normalizes in larger tumors at later timepoints. (*A*) Representative H&E-stained sections of tumors dissected from parental and mutant teratomas at day 28. Regions surrounded by black lines highlight immature neuroepithelial elements. Blue lines highlight mature epithelium (squamous, respiratory, and/or gastroinstestinal). Red asterisk represents mature neural regions. Green lines show mature cartilage. Yellow lines show mature fat. Orange lines show yolk sac (immature). (*B*) Average percent immature component for day 28 parental homozygous mutant teratomas. N=3 individual tumors. Error bars denote ±standard deviation. (*C*) Similar analysis performed for a different parent and homozygous Oct4$^{C48S}$ mutant BL/6 ESC pair grown as teratomas for 28 days. N=5.

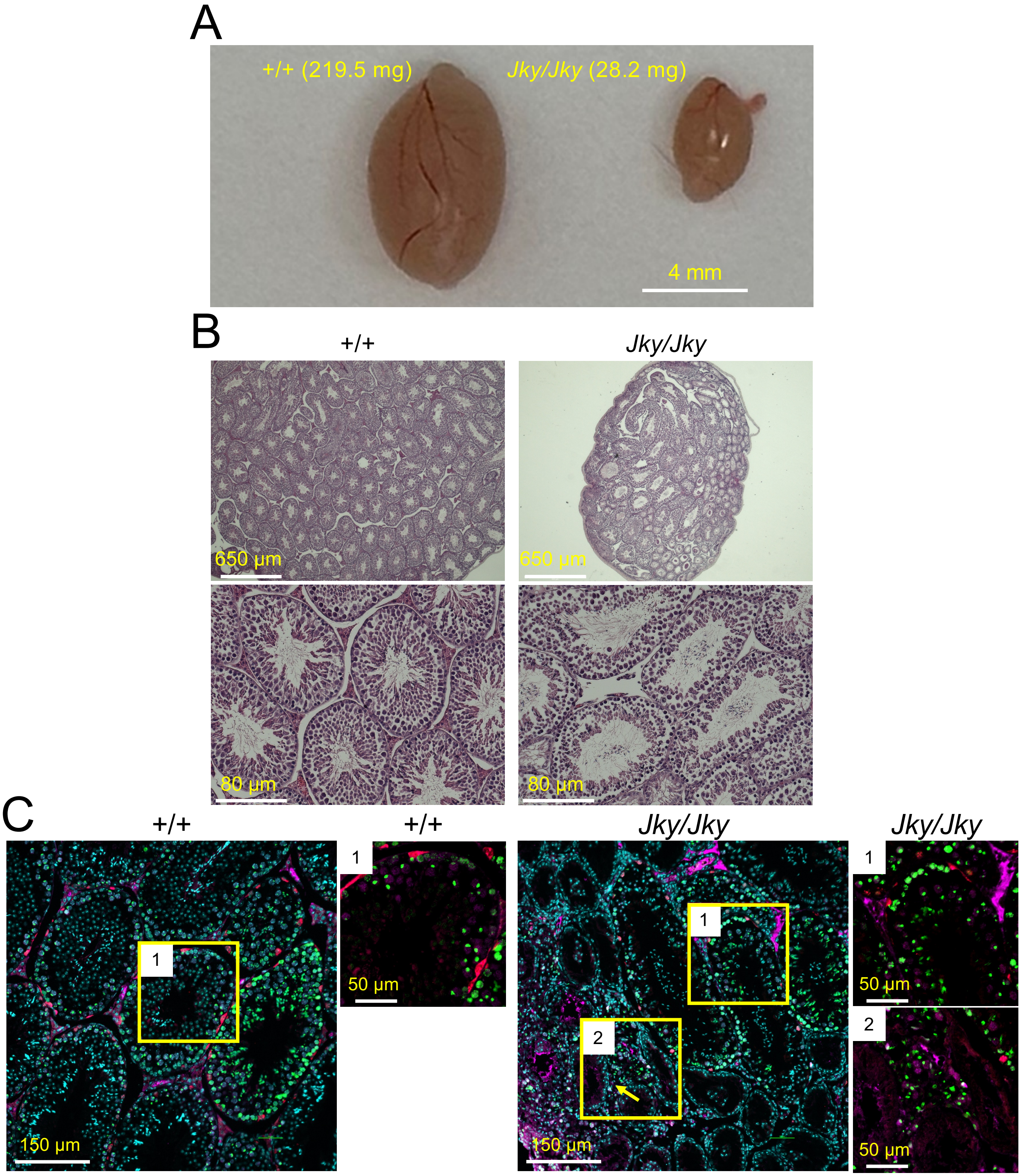


Cyan: DAPI Red: LIN28A (spermatogonia) Green: pH2AX (DNA damage and pachytene spermatocyte) Violet: SYCP3 (spermatocyte)

**Figure S13.** Abnormal testes in *Jky* homozygous mice. (*A*) Gross images of dissected testes from a 2.5 month-old male *Jky*/*Jky* mouse and +/+ littermate control. Averaged weights of two testes from each animal are also shown. (*B*) H&E staining of FFPE testes sections from *Jky*/*Jky* and +/+ littermate control testes. (*C*) Representative immunofluorescence images showing abnormal spermatogenesis and germ cell loss in testis sections from the *Oct4*$^{Jky/Jky}$ mouse: gamma H2AX (green); LIN28A (Red); SYCP3 (violet); and DNA (Cyan). Box #1 highlights examples of superficially normal spermatogenesis, while #2 highlights example abnormal localization of spermatocytes and germ cell loss. Arrow points to abnormal localization of spermatocytes within interstitial regions.

Fig.1C:

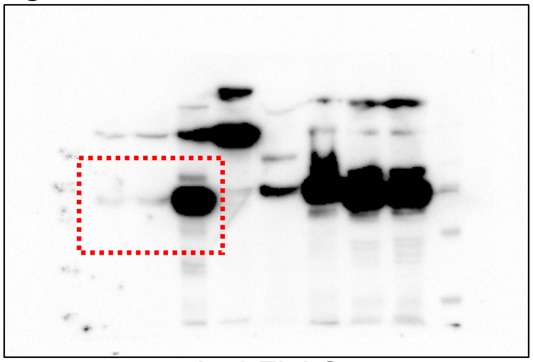

Anti-FLAG

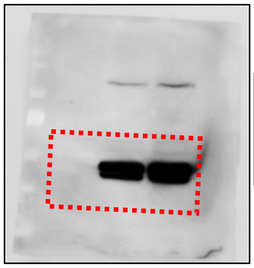

Anti-Oct4

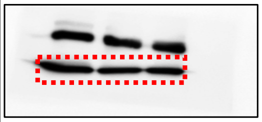

Anti-H3

Fig.1F:

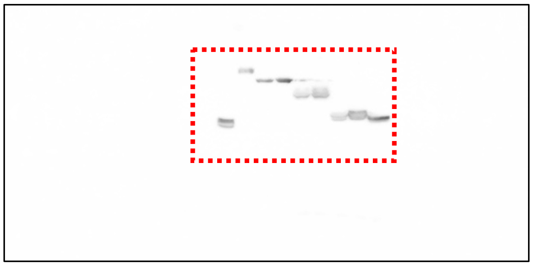

Anti-FLAG

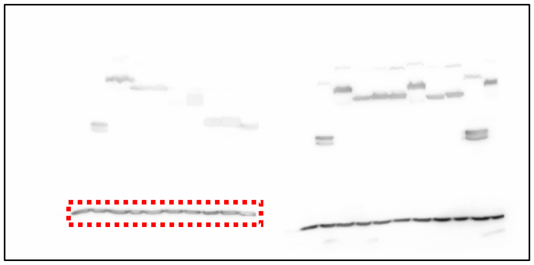

Anti-H3

Fig.2C:

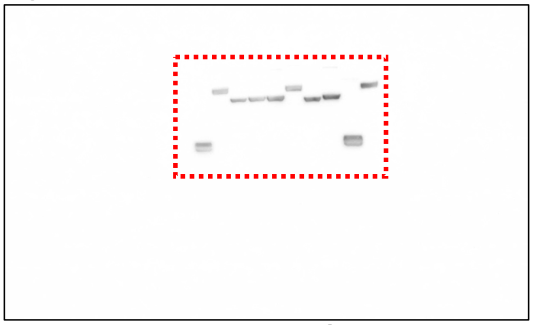

Anti-FLAG

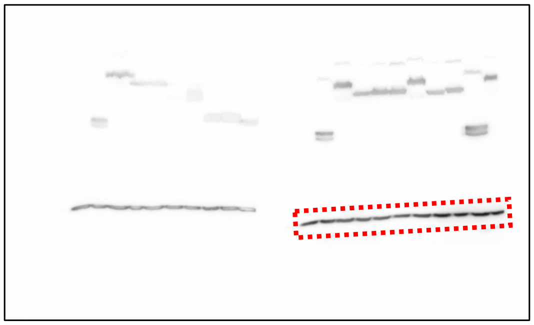

Anti-H3

Fig.3E:

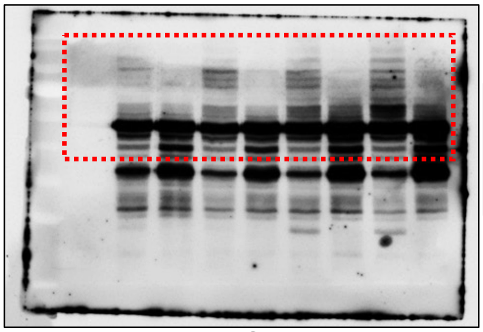

Anti-Oct4

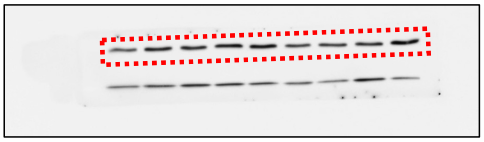

Anti-H3

Fig.3F:

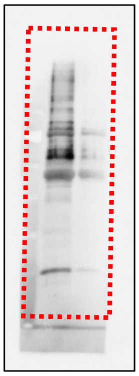

Anti-FK2

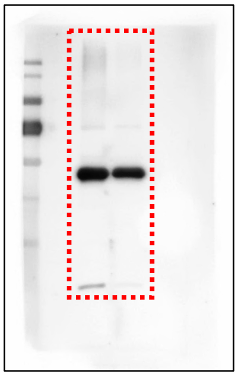

Anti-Ub-K48

Fig.3G:

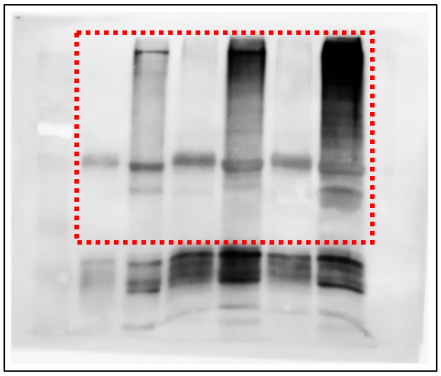

Anti-FK2

**Figure S14.** Uncropped immunoblots for Figs. 1-3.

Shen et al. Figure S15

Fig.4B:

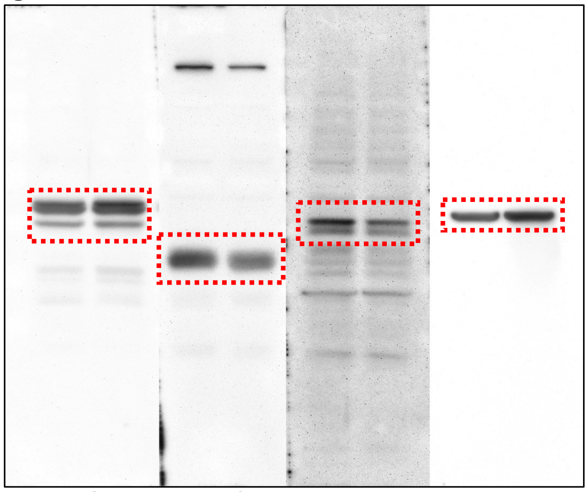

Anti-Oct4 Anti-Sox2 Anti-Nanog Anti-β-actin

Fig.4D:

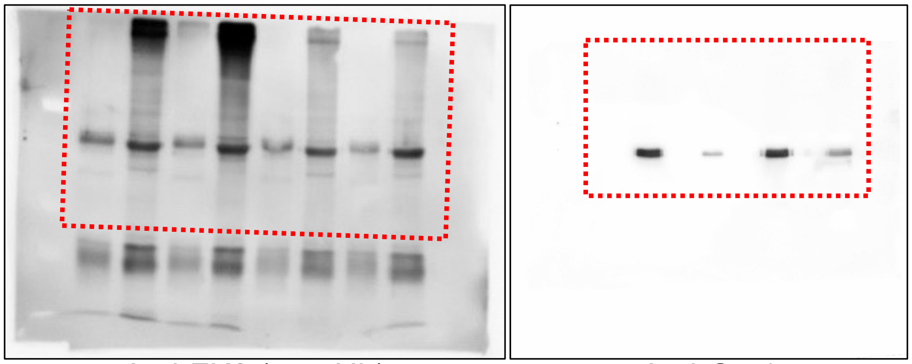

Anti-FK2 (pan-Ub) Anti-Oct4

Fig.4E:

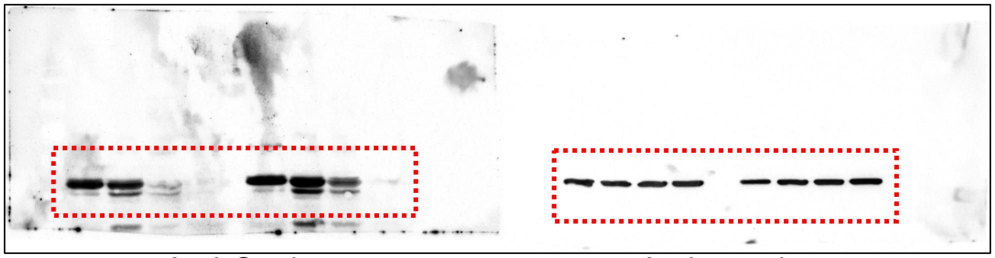

Anti-Oct4 Anti-β-actin

**Figure S15.** Uncropped immunoblots for Fig. 4.

Shen et al. Figure S16

Supp. Fig.S1B:

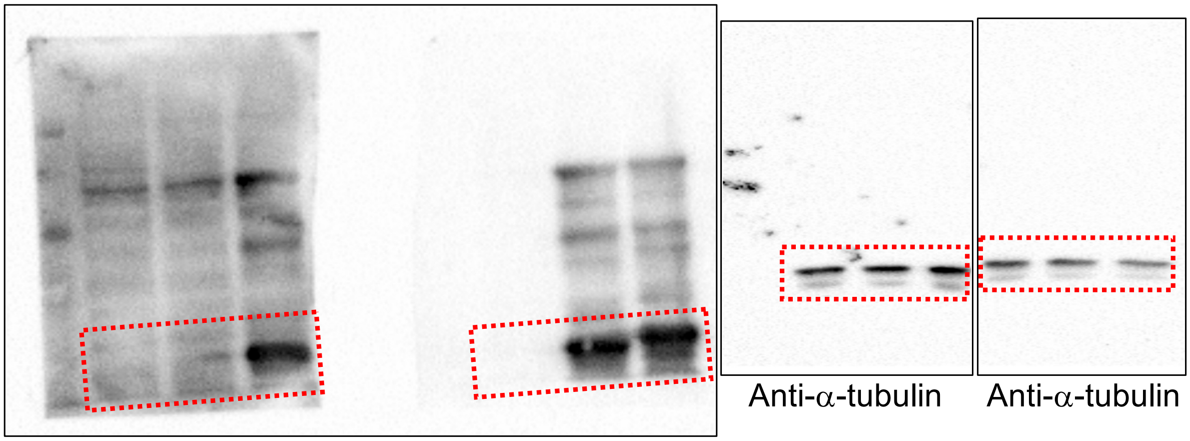

Anti-α-tubulin Anti-α-tubulin

Anti-FLAG

Supp. Fig.S8:

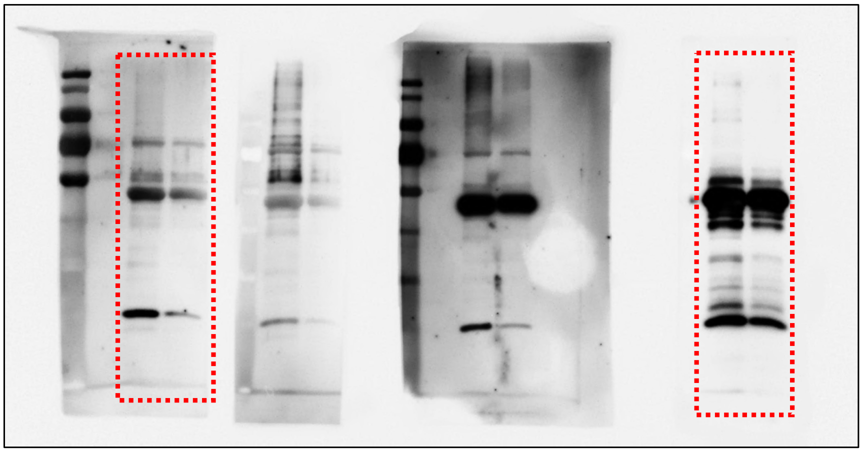

Anti-Ub-K63 Anti-Oct4

**Figure S16.** Uncropped immunoblots for Suppl. Fig. S1 and S8.

### Supp. Fig.S9C:

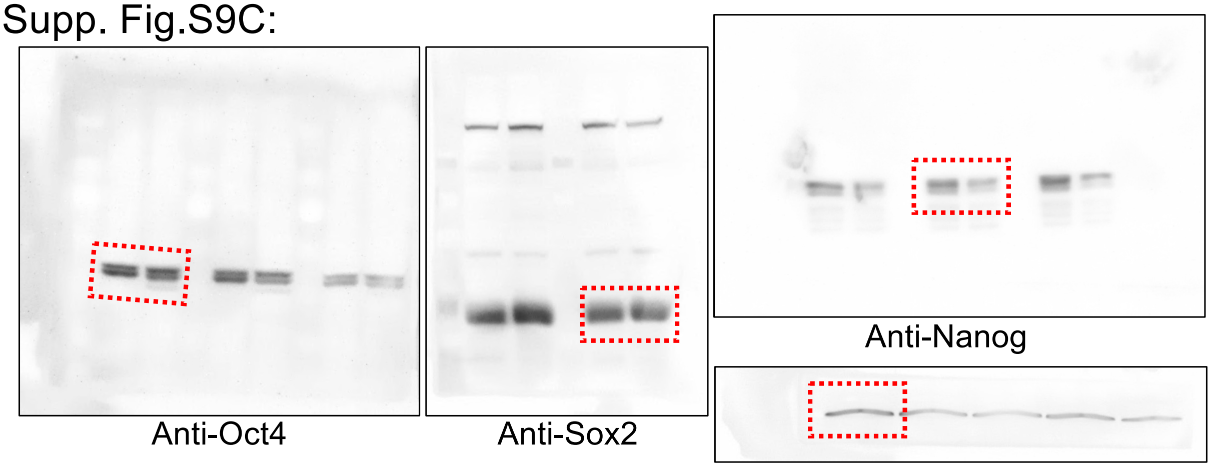


### Supp. Fig.S9E:

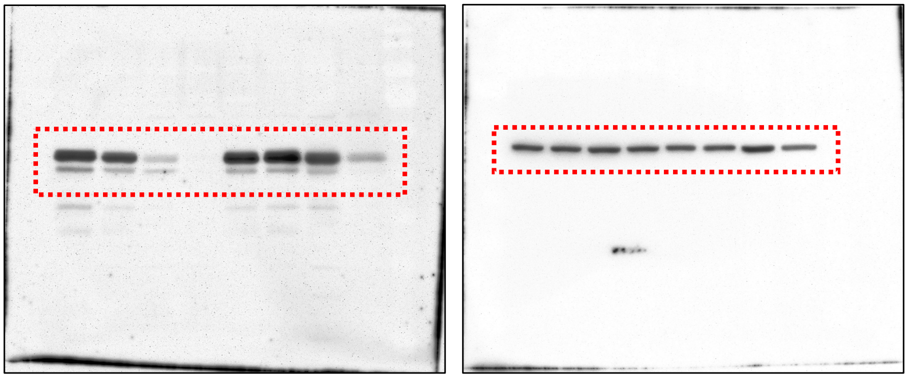


### Supp. Fig.S9F:

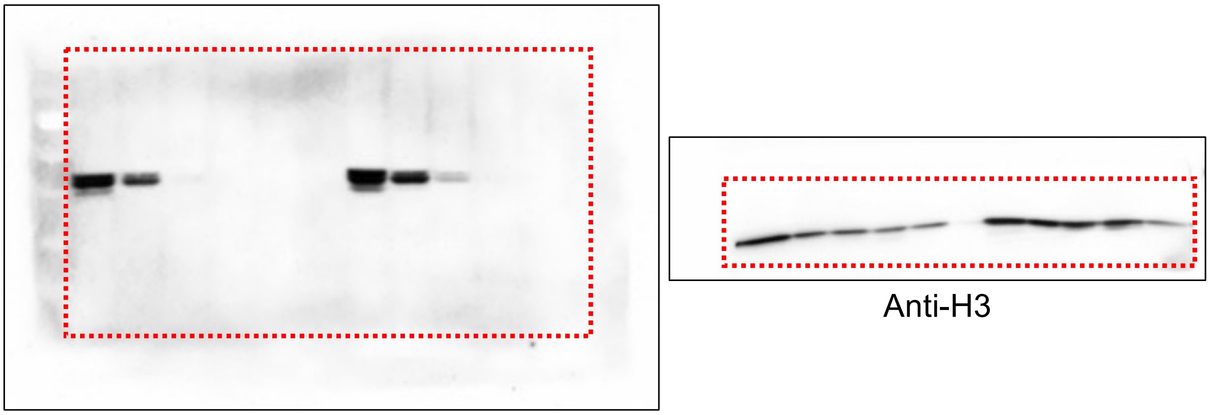


### Supp. Fig.S9G:

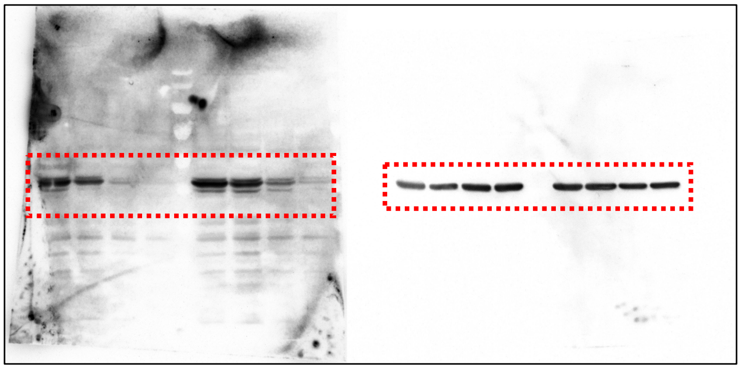

**Figure S17.** Uncropped immunoblots for Suppl. Fig. S9.
